# Evidence of transcription at polyT short tandem repeats

**DOI:** 10.1101/634261

**Authors:** Chloé Bessière, Manu Saraswat, Mathys Grapotte, Christophe Menichelli, Jordan A. Ramilowski, Jessica Severin, Yoshihide Hayashizaki, Masayoshi Itoh, Akira Hasegawa, Harukazu Suzuki, FANTOM consortium, Piero Carninci, Michiel J.L. de Hoon, Wyeth W. Wasserman, Laurent Bréhélin, Charles-Henri Lecellier

## Abstract

**Background:** Using the Cap Analysis of Gene Expression technology, the FANTOM5 consortium provided one of the most comprehensive maps of Transcription Start Sites (TSSs) in several species. Strikingly, ~72% of them could not be assigned to a specific gene and initiate at unconventional regions, outside promoters or enhancers.

**Results:** Here, we probe these unassigned TSSs and show that, in all species studied, a significant fraction of CAGE peaks initiate at short tandem repeats (STRs) corresponding to homopolymers of thymidines (T). Additional analyses confirm that these CAGEs are truly associated with transcriptionally active chromatin marks. Furthermore, we train a sequence-based deep learning model able to predict CAGE signal at T STRs with high accuracy (~81%) Extracting features learned by this model reveals that transcription at T STRs is mostly directed by STR length but also instructions lying in the downstream sequence. Excitingly, our model also predicts that genetic variants linked to human diseases affect this STR-associated transcription.

**Conclusions:** Together, our results extend the repertoire of non-coding transcription associated with DNA tandem repeats and complexify STR polymorphism. We also provide a new metric that can be considered in future studies of STR-related complex traits.

## Background

RNA polymerase II (RNAP-II) transcribes many loci outside annotated protein-coding gene (PCG) promoters [1, 2] to generate a diversity of RNAs, including for instance enhancer RNAs [3] and long non-coding RNAs [4]. In fact, *>* 70% of all nucleotides are thought to be transcribed at some point [1, 5, 6]. Non-coding transcription is far from being fully understood [7] and some authors suggest that many of these transcripts, often faintly expressed, can simply be ‘noise’ or ‘junk’ [8]. On the other hand, many non annotated RNAP-II transcribed regions correspond to open chromatin [1] and *cis*-regulatory modules (CRMs) bound by transcription factors (TFs) [9]. Besides, genome-wide association studies showed that trait-associated loci, including those linked to human diseases, can be found outside canonical gene regions [10–12]. Together, these findings suggest that the non-coding regions of the human genome harbor a plethora of potentially transcribed functional elements, which can drastically impact genome regulations and functions [12, 13]. Notably, short tandem repeats (STRs), repeated DNA motifs of 2 to 6 bp, constitute one of the most polymorphic and abundant repetitive elements [14]. STR polymorphism, which corresponds to variation in number of repeated DNA motif (i.e. STR length), is due to their susceptibility to slippage events during DNA replication. STRs have been shown to widely impact gene expression and to contribute to expression variation [15, 16]. At the molecular level, they can affect transcription by inducing inhibitory DNA structures [17] and/or by modulating the binding of transcription factors [18, 19]. Using the Cap Analysis of Gene Expression (CAGE) technology [20, 21], the FANTOM5 consortium provided one of the most comprehensive maps of TSSs in several species [2]. Integrating multiple collections of transcript models with FANTOM CAGE datasets, Hon *et al.* built an atlas of 27,919 human lncRNAs, among them 19,175 potentially functional RNAs, and provided a new annotation of the human genome (FANTOM5 CAGE Associated Transcriptome, FANTOM CAT) [4]. Despite this annotation, many CAGE peaks remain unassigned to a specific gene and/or initiate at unconventional regions, outside promoters or enhancers, providing an unprecedented mean to further characterize non-coding transcription within the genome ‘dark matter’ [13] and to decode part of the transcriptional noise.

Here, we probed CAGE data collected by the FANTOM5 consortium [2] and specifically looked for repeating DNA motifs around CAGE peak summits. In all species studied, we showed that a fraction of CAGE peaks (between 2.22% in rat and 6.45% in macaque) initiate at repeats of thymidines (Ts) i.e. STRs of Ts (or T STRs). Biochemical and genetic evidence demonstrate that many of these CAGEs do not correspond to technical artifacts, as previously suspected [22], but rather exhibit genuine features of TSSs. Our results not only extend the repertoire of non-coding transcription but also complexify T STR polymorphism, with T STR having the same length but different transcription rate and, conversely, T STR with different lengths having similar transcription rate. We further learned a sequence-based Convolutional Neural Network (CNN) able to predict this transcription with high accuracy (∼ 81% in human). Extracting the features learned by this model reveals that this transcription is triggered by the length of T STRs but also by instructions lying in the downstream sequence. Using our CNN model, we finally showed that many genetic variants linked to human diseases potentially affect this STR-associated transcription, thereby advancing our capacity to interpret several regulatory variants [15].

## Results

### A significant fraction of CAGE peaks initiates at T homopolymers in various species

Using the motif enrichment tool HOMER [23], we looked at repeating DNA motifs in 21bp-long sequences centered around the FANTOM5 CAGE peak summits. As shown in Figure 1, the first motif identified in both human and mouse is the canonical initiator element INR [24, 25], demonstrating the relevance of our strategy to unveil specific sequence-level and TSS-associated features. A second motif corresponding to T homopolymer is identified (Figure 1A and B). This motif is present in 61,907 human and 8,274 mouse CAGE peaks (Figure 1A and C). Because homopolymers of Ts represent the most abundant class of STRs [14, 26], we hereafter called these CAGEs, T STR CAGEs. In human, the median size of CAGE-associated T STRs is 17 bp with a minimum of 9 bp and a maximum size of 64 bp. Similar results are obtained in mouse (median = 21 bp, minimum = 8 bp and maximum = 58 bp). Looking directly at the number of STRs of ≥ 9 Ts in the human genome, we found that 63,974 T STRs out of 1,337,561 exhibit a CAGE peak located at their 3’end + 2bp. Thus, the vast majority of T STR CAGEs are identified by motif enrichment and not all T STRs are associated with CAGE peaks. In mouse, only 8,825 T STRs out of 834,954 are associated with a CAGE peak but, compared to human CAGE data, mouse data are small-scaled in terms of number of reads mapped and diversity in CAGE libraries [2]. We further looked at CAGE peaks in dog, chicken, macaque and rat (Supplementary Figure S1) and T homopolymer is invariably detected by HOMER (sometimes even before INR motif).

**Figure 1.**
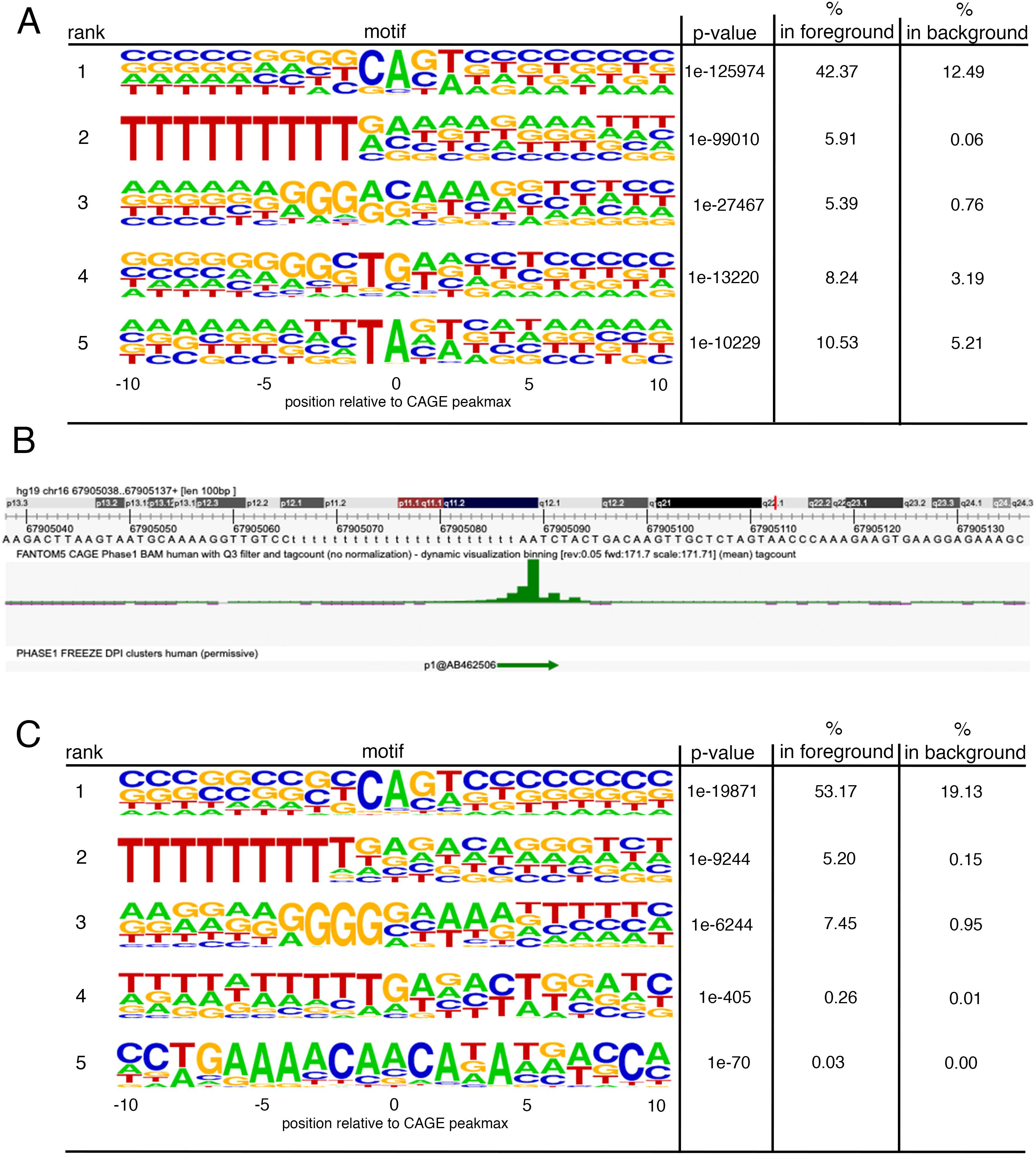
Motif discovery around CAGE peakmax. HOMER [23] was used to find 21bp-long motifs in 21bp-long sequences centered around the peak summit of FANTOM5 CAGEs. A. 1,048,124 human permissive CAGEs were used. The top 5 is shown. 61,907 CAGEs are associated with T homopolymers starting at −2 (0 being the peakmax). Motif enrichment is indicated by binomial test p-value. B. Example of T homopolymer located at chr16:67905066 and associated with a CAGE peak (p1@AB462506). Upper track shows the DNA sequence. Middle track shows the CAGE signal averaged in 988 human FANTOM libraries. Bottom track shows the CAGE peak calling as defined in [2]. C. Among 158,966 mouse CAGEs, 8,274 are associated with T homopolymers. Note that another repeat of 2-5 Gs is also identified in FANTOM CAGE data with strong similarities in both human and mouse (A and C). However, this repeat is less strict and is not conserved in dog, chicken, macaque and rat (Supplementary Figure S1) and is absent in Start-seq and DECAP-seq data (Supplementary Figure S3). Though this motif may represent a biologically relevant signal in human and mouse, these features make it less relevant for our study.

The position of the CAGE summit at −2 is presumably artificially introduced by the hybridization step on the flow cell (Supplementary Figure S2A): sequencing of any CAGE tags initiating within T homopolymers will preferentially start just after polyT track. The motif enrichment is therefore more indicative of transcription initiation at T STRs than of the existence of a true motif located precisely 2bp after T STRs. In line with this, ENCODE CAGE data, which were generated using Illumina technology, confirmed several FANTOM CAGE peaks (generated by HeliscopeCAGE) but were not exactly aligned on that of FANTOM (Supplementary Figure S2B). Note that the FANTOM CAT annotation was actually shown to be more accurate in 5’ end transcript definitions compared to other catalog [4]. Moreover, a small fraction of Start-seq [27] and DECAP-seq [28] TSSs can also initiate in sequences with a preference for T and this clearly does not represent a genuine motif (Supplementary Figure S3A and B). A similar observation was made with TSS-seq data collected in *Arabidopsis thaliana* [29] (Supplementary Figure S4, motif #2).

Provided that Heliscope sequencing used by FANTOM5 can be internally primed at polyT track (Supplementary Figure S2), we can legitimately question the relevance of T STR CAGEs [22]. Several features are indeed in agreement with the idea that T STR CAGEs could for instance arise from internal priming within introns of messenger RNAs (Supplementary Figures S5 to S9 and Supplementary Tables S1 and S2). However, this artefact scenario could not explain all T STR CAGEs as 8,926 (out of 63,974, *>* 14%) are ‘intergenic’, i.e. not located in the same orientation as one of the 53,220 genes of the FANTOM CAT annotation, one of the largest gene annotations so far. No major sequence difference distinguishes intergenic from intragenic T STR CAGEs (Supplementary Figure S10). Besides, we observed some concordance between several technologies, which do not use oligo dT priming (Supplementary Figures S2B, S3, S4 and S9). Together these observations raise the possibility that a fraction of T STR CAGEs represent genuine TSSs. The corresponding RNAs appear rather stable [30] according to the CAGE exosome sensitivity score previously computed by FANTOM [4] (median sensitivity score = 0.08, Supplementary Figure S6B), suggesting that they do not correspond to cryptic transcription [7]. To clarify the existence of these TSSs, we further investigated whether T STR CAGEs exhibit canonical TSS features.

### Several T STR CAGE tags are truly capped

We used a strategy described by de Rie *et al.* [31], which compares CAGE sequencing data obtained by Illumina (ENCODE) vs. Heliscope (FANTOM) technologies. Briefly, the 7-methylguanosine cap at the 5’ end of CAGE tags produced by RNA polymerase II can be recognized as a guanine nucleotide during reverse transcription. This artificially introduces mismatched Gs at Illumina tag 5’ end, which is not detected with Heliscope [31]. Although such G bias is not clearly observed when considering the whole CAGE peak, it is readily detectable when considering the 3’ end of T STRs (position −2 from FANTOM CAGE summit) using Illumina ENCODE CAGE data produced in Hela-S3 nuclei (Figure 2A). This bias is observed in other cell types and is comparable to that observed with CAGE tags assigned to gene TSSs (Figure 2B and Supplementary Figure S11). The G bias is even more pronounced in intergenic than intragenic T STR CAGEs, suggesting that reads corresponding to host gene mRNA can mask transcription at STRs (Figure 2B and Supplementary Figure S11). Conversely, most CAGE tag 5’ ends perfectly match the sequences of pre-miRNA 3’end, as previously reported [31], or that of 61,907 randomly chosen genomic positions (Figure 2B and Supplementary Figure S11). Mismatched Gs at the 3’ end of all T STRs located within the same genes as T STR CAGEs are also detected (Figure 2B and Supplementary Figure S11), though the abundance of tags is higher at T STR CAGEs (Figure 2B), suggesting the potential existence of false negatives (see below). Overall, these analyses show that T STR CAGEs are truly capped as opposed to post-transcriptional cleavage products of pri-mRNAs [31] or to random genomic positions.

**Figure 2.**
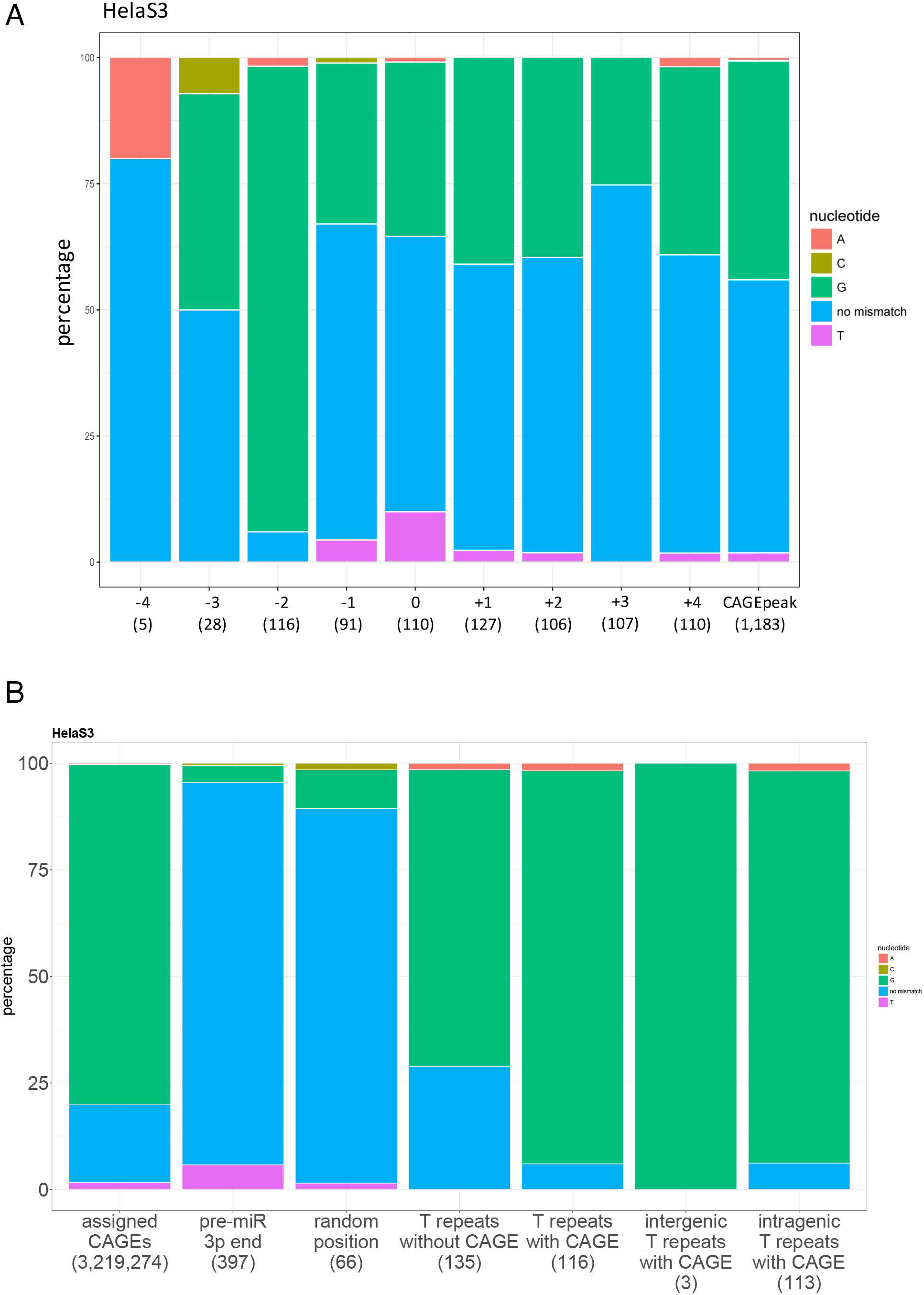
T STR CAGEs are 5’-capped. A. G bias in ENCODE CAGE reads (nuclear fraction, polyA-) was assessed at the indicated positions (x-axis) around T STR CAGEs (0 corresponds to end of poly repeat + 2bp, n = 63,974). The number of intersecting reads on the positive strand is indicated in bracket. B. Same analyses as in A but considering position −2 only and other CAGEs/positions indicated in x-axis: CAGE peaks assigned to gene, pre-microRNA 3’ ends, 61,907 random positions, T repeat without CAGE. Note that the number of T STR CAGEs on + strand is 33,209 and that of T STRs without CAGE 636,993 indicating that the number of reads at T STRs with CAGE is relatively higher than that at T STRs without CAGE. Intergenic (not located in, and in the same strand of, one of the FANTOM CAT genes) and intragenic (located in, and in the same strand of, one of the FANTOM CAT genes) T STR CAGEs were also distinguished. Color code as in A.

### Many T STR CAGEs are associated with transcription-related epigenetic marks

We then determined whether T STR CAGEs are associated with epigenetics marks related to transcription at the chromatin level. Using ENCODE DHS Master list, we confirmed that ∼ 11% of T STR CAGEs (7,028 out of 63,974) lie in DHSs, while this is true for only ∼ 6% of T STRs without CAGE peaks (76,616 out of 1,273,587, Fisher’s exact test p-value *<* 2.2e-16). This difference was also found with another compendium of DHSs (https://www.meuleman.org/project/dhsindex/: 7,028 out of 63,974 T STRs with CAGE peaks were found in DHSs while only 76,616 out of 1,273,587 T STRs without CAGE peaks, Fisher’s exact test p-value *<* 2.2e-16) as well as with ATAC-seq data [32] (12% of T STR CAGEs (7,697 out of 63,974) but only 6.6% of T STRs without CAGE peaks (84,459 out of 1,273,587) were found in pan-cancer open regions, Fisher’s exact test p-value *<* 2.2e-16).

Since most T STR CAGEs are intronic, the differences observed may merely be due to the transcriptional state of the host gene and/or the genomic environment. We then investigated transcription at T STRs at the gene level and library-wise in order to preclude the effect of global chromatin/gene environment. We created two sets of ‘expressed’ and ‘non-expressed’ T STR CAGEs: T STR CAGEs are considered as ‘expressed’ if (i) associated with a detectable CAGE signal in the sample considered (TPM *>* 0) and (ii) located in a gene containing at least one ‘non-expressed’ T STR CAGEs. Conversely, ‘non-expressed’ T STR CAGEs are (i) not detected in the sample considered but detected in other samples and (ii) located in a gene containing at least one ‘expressed’ T STR CAGEs. Using these two sets of T STR CAGEs makes the analyses independent of host gene expression because the same genes are considered in both cases. Genome segmentation provided by combined ChromHMM and Segway [33, 34] shows that ‘expressed’ T STR CAGEs are systematically more enriched in regions corresponding to predicted transcribed regions than ‘non expressed’ T STR CAGEs (Figure 3A, Fisher’s exact test p-value *<* 2.2e-16 in HeLa-S3 (CNhs12325) and in GM12878 (CNhs12331), p-value = 2.053e-13 in K562 (CNhs12334)). The ChromHMM/Segway segmentations integrate several ChIP-seq data corresponding to RNA polymerase II, 8 chromatin marks (H3K4me1, H3K4me2, H3K4me3, H3K27ac, H3K9ac, H3K36me3, H4K20me1, H3K27me3) and the CTCF transcription factor. Using the same comparison, transcription at T STRs was also confirmed with GRO-seq data (Figure 3B and Supplementary Figures S12 and S13).

**Figure 3.**
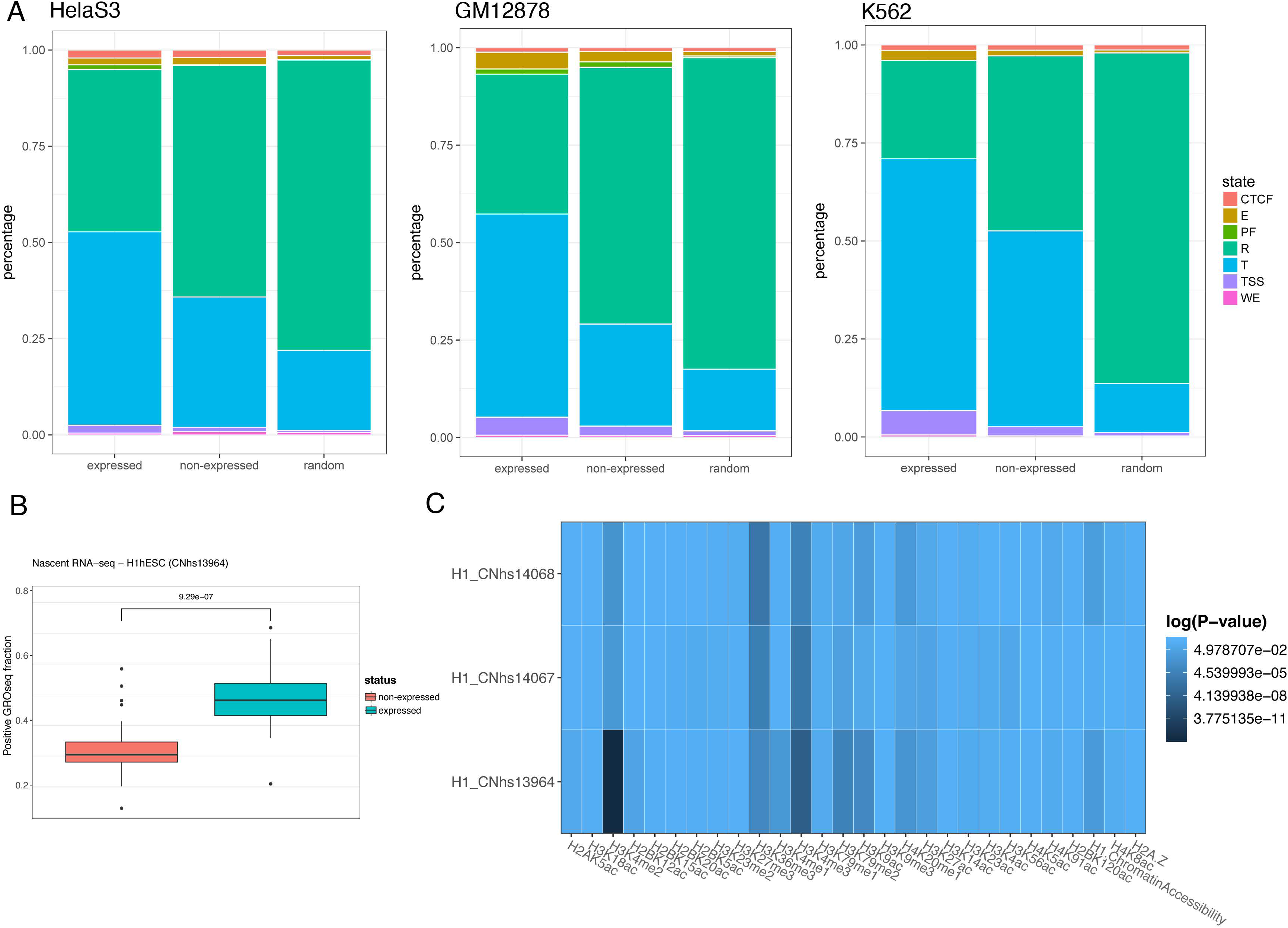
Several T STR CAGEs are associated with promoter-related epigenetics marks. **A.** Distribution of ChromHMM/Segway genome segments containing ‘expressed’ or ‘non-expressed’ T STR CAGEs (see text for definitions) in HeLa-S3, GM12878 and K562. 61,907 random position were also considered for comparison. TSS, Predicted promoter region including TSS; PF, Predicted promoter flanking region; E, Predicted enhancer; WE, Predicted weak enhancer or open chromatin cis regulatory element; CTCF, CTCF enriched element; T, Predicted transcribed region; R, Predicted Repressed or Low Activity region. Note the absence of ‘Predicted promoter flanking region’ class in K562. **B.** Fraction of T STR CAGEs ‘expressed’ (blue) or not (red) in H1 embryonic stem cells (CNhs13964 FANTOM library) and intersecting with nascent RNAseq signal from 26 ‘gro-seq colorado’ H1-ESC samples (considering a window of 101bp centered around T repeat end). Boxplots show the results of the intersection for 26 samples (see Supplementary Figures S12 and S13 for separated analyses). Fisher’s exact test indicates that the fraction of T STR CAGEs with no-null GROseq signal obtained for ‘expressed’ T STR CAGEs is higher than that obtained for ‘non expressed’ T STR CAGEs (p-value *<* 2.2e-16). **C.** Intersection of ‘expressed’/’non expressed’ T STR CAGEs coordinates with that of Roadmap epigenetics data collected in H1 embryonic stem cells. The fraction obtained for ‘expressed’ and ‘non expressed’ T STR CAGEs were compared using Fisher’s exact test. The color correspond to p-value *<* 5e-3 (green) or ≥ 5e-3 (grey).

We then looked at epigenetics marks individually and used data collected by the Roadmap Epigenome consortium in H1 embryonic stem cells to compare the epigenetics status of ‘expressed’ vs. ‘non-expressed’ T STR CAGEs. In three replicates of untreated H1 cells CAGE libraries, H3K36me3 and H3K4me3 peaks are invariably enriched in ‘expressed’ T STR CAGEs (Figure 3C and Supplementary Figure S14). Similar profiles are obtained with ENCODE ChIP-seq data, although less pronounced in GM12878 (Supplementary Figure S15). H3K4me3 levels at T STR CAGEs are low (Supplementary Figures S14 and S15), as observed with lncRNAs [35]. H3K36me3 is a histone modification mark enriched in the gene body region and associated with transcription elongation [35]. H3K4me3 is a mark classically associated with active or poised transcription start sites [35]. Hence, ‘expressed’ T STR CAGEs are more associated with H3K4me3/H3K36me3 domains than ‘non-expressed’ ones. Interestingly, this type of ‘K4-K36’ domains have been previously used to characterize lncRNAs [36]. Figure 3C also shows an enrichment in H3K4me2 (see also Supplementary Figure S14 and S15), which was previously associated with intragenic *cis*-regulatory [37] and TF binding [38] regions. We concluded that, overall, detection of CAGEs at T STRs is associated with transcription-related chromatin marks.

### Several T STR CAGEs correspond to annotated transcript and enhancer TSSs

We found that 11,180 T STRs (end+2bp) were associated with a ‘robust’ CAGE peak i.e. peaks confirmed by external data of ESTs and associated with H3K4me3 marks and DNase hypersensitivity sites (DHSs) [2], representing *>* 17% of all T STR CAGEs, and thereby confirming the results shown in Figure 3. We then assessed the presence of T STR transcription among annotated TSSs. Motif enrichment around FANTOM CAT TSSs [4] shows that ∼ 1.5% of them initiate downstream T STRs (Figure 4A). Looking directly at TSS coordinates of FANTOM CAT robust transcripts [4], we noticed that 6,734 TSSs, corresponding to 10,606 robust transcripts (out of 525,237, ∼ 2%), initiate 2bp after a T STR, with a clear enrichment in lncRNA intergenic, lncRNA sense intronic or sense overlap RNA (hypergeometric test p-values *<* 2.2e-16 for the 3 RNA classes). A list of these TSSs is provided in Supplementary Table S3A. Similar results were obtained for 5,889 stringent transcripts (total n = 402,813) [4]. Importantly, transcript models in FANTOM CAT combine various independent sources (GENCODE release 19, Human BodyMap 2.0, miTranscriptome, ENCODE and an RNA-seq assembly from 70 FANTOM5 samples) and FANTOM CAT TSSs were validated with Roadmap Epigenome DHS and RAMPAGE data sets [4]. As expected, these TSSs only moderately contribute to gene expression (Supplementary Figure S16). No specific Open Reading Frame could be detected 2kb downstream T STR CAGEs (Supplementary Figure S17).

**Figure 4.**
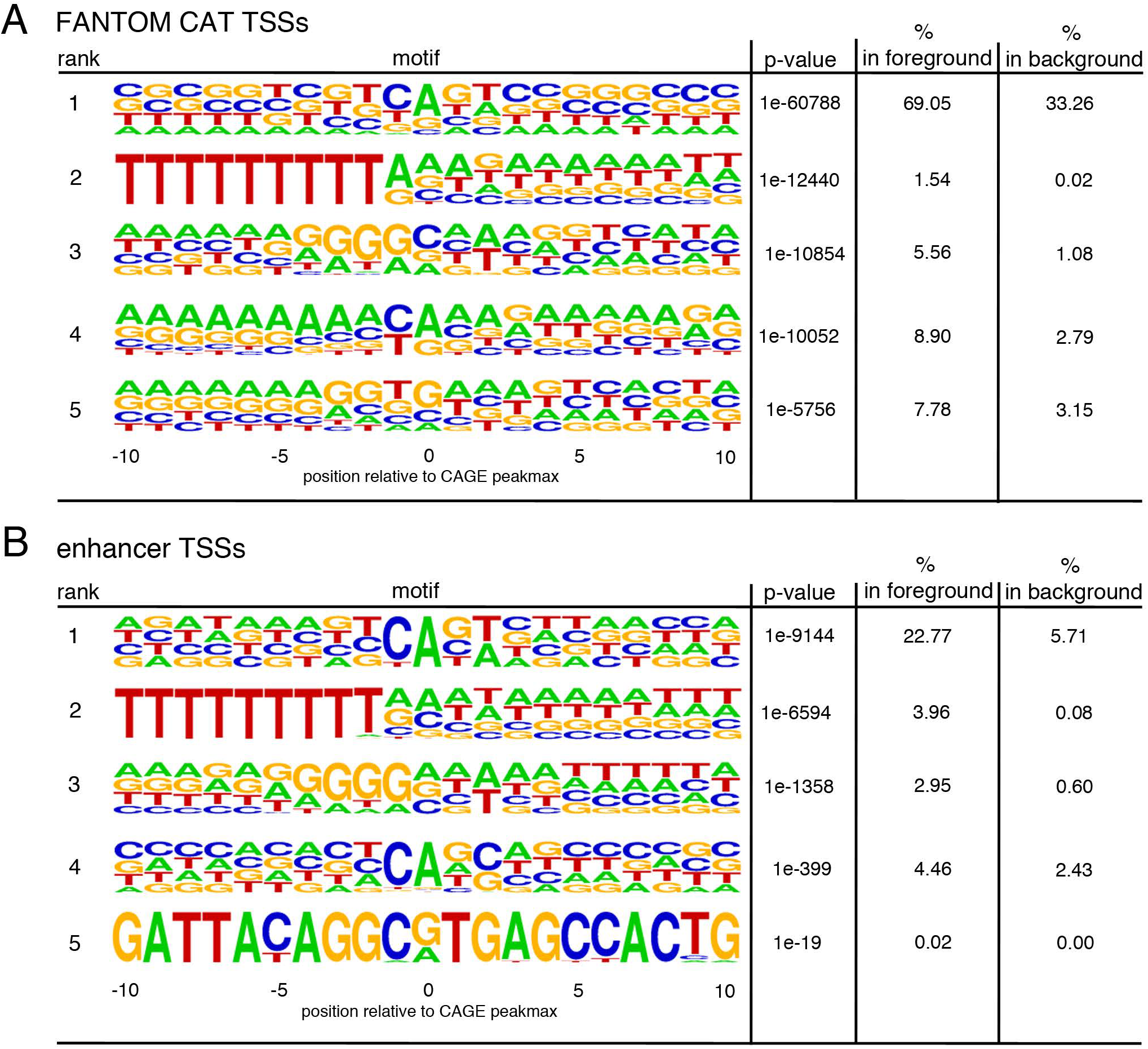
Motif discovery around FANTOM CAT transcript and enhancer TSSs. HOMER [23] was used to find 21bp-long motifs in 21bp-long sequences centered around TSSs of FANTOM CAT robust transcripts (A) or enhancers (B). A. 525,237 transcript starts as defined in [4] were used. The top 5 is shown. Motif enrichment is indicated by binomial test p-value. B. 65,423 enhancers defined in [3] were used corresponding to 130,846 TSSs. Only 5 motifs were identified by HOMER.

The T homopolymer is also observed for ∼ 4% of FANTOM enhancer TSSs [3] (Figure 4B), though no enrichment in enhancer epigenetic marks are predicted by chromHMM/Segway (Figure 3A). Looking directly at genomic coordinates, 4,976 enhancers (out of 65,423, ∼ 7.6%) are defined with at least one T STR CAGE (Supplementary Table S3B) and 173 enhancers are defined by two T STR CAGEs [3]. Enhancer TSSs are therefore more often associated with T STRs than transcript TSSs (∼ 7.6% and ∼ 2% respectively, Fisher’s exact test p-value *<* 2.2e-16). Similar results were obtained with mouse enhancers (Supplementary Figure S18) with 1,171 enhancers (out of 44,459, ∼ 2.6%) involving at least one T STR CAGE. These results strengthen the idea that a number of T STR CAGEs correspond to genuine TSSs.

### T STR CAGEs interact with gene TSSs

We used ENCODE Chromatin Interaction Analysis by Paired-End Tag Sequencing (ChIA-PET) directed against RNAP-II in K562 cell line [39] to better characterize the genetic elements interacting with T STR CAGEs. As in Figure 3, we used the sets of ‘expressed’ and ‘non-expressed’ T STR CAGEs in K562 cells. The ‘expressed’ T STR CAGEs are slightly enriched in ChIA-PET data compared to ‘non-expressed’ T STR CAGEs (Figure 5A), confirming the results depicted in Figure 3. Using combined chromHMM/Segway, we observed that regions interacting with ‘non-expressed’ T STR CAGEs are enriched in ‘Predicted Repressed or Low Activity’ regions (Figure 5B and Supplementary Figure S19). On the other hand, regions interacting with ‘expressed’ T STR CAGEs are enriched in ‘Predicted promoter region including TSS’ regions (Figure 5B and Supplementary Figure S19). We further used the FANTOM5 CAGE TSS classification [2] and showed that CAGEs interacting with ‘expressed’ T STR CAGEs preferentially interact with ‘strict’ and ‘weak’ TSSs, compared to ‘non-expressed’ T STR CAGEs (Figure 5C and Supplementary Figure S20). Note that ChIA-PET data contain too many intragenic interactions to clearly conclude that these TSSs correspond to host gene TSSs.

**Figure 5.**
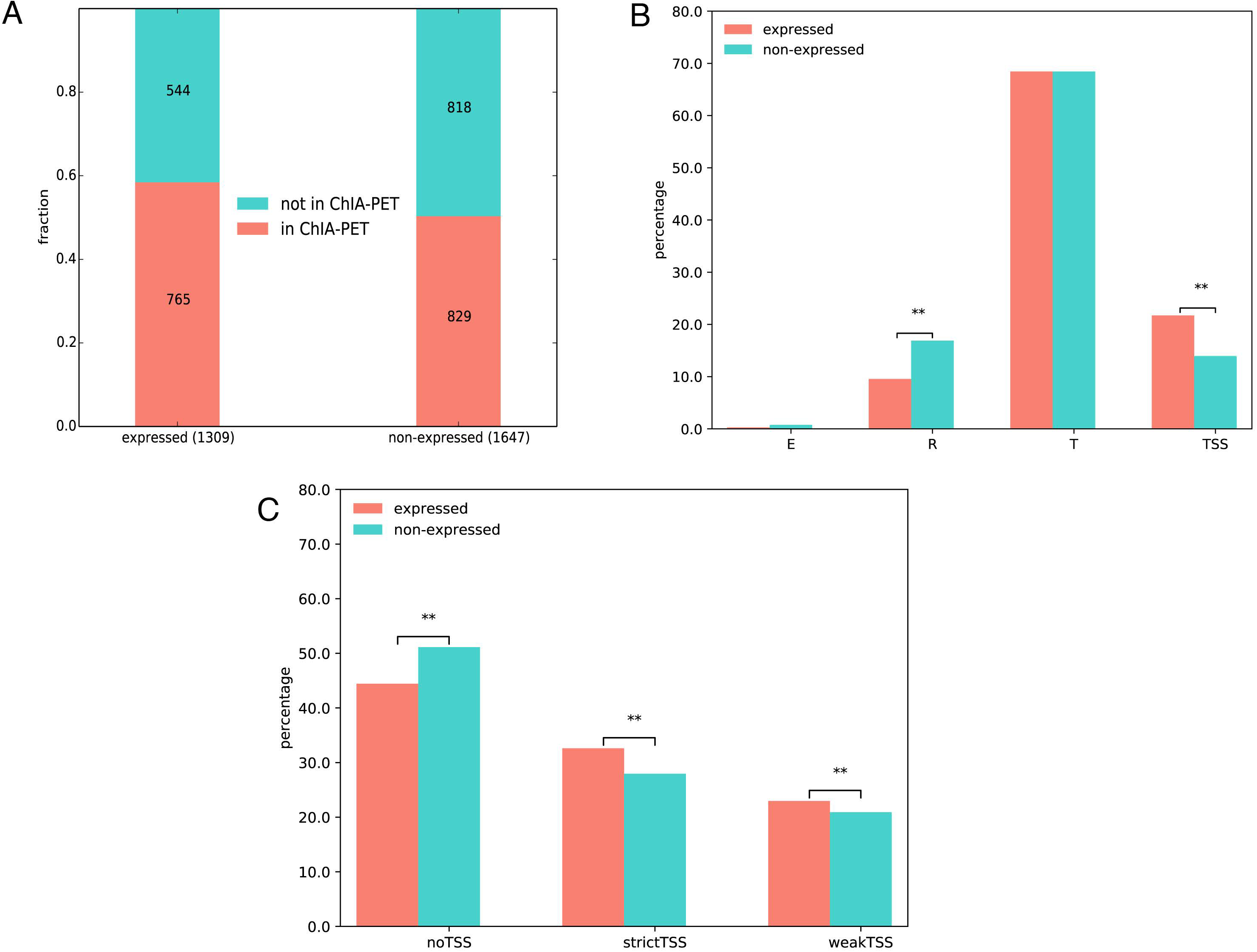
T STR CAGEs interact with gene TSSs. **A.** The number of T STR CAGEs associated with a CAGE signal (’expressed’) or not (’non-expressed’) in K562 cells located (red) or not (green) in ChIA-PET interacting regions was calculated intersecting their genomic coordinates. Results are shown in the case of CNhs12334 CAGE and RNAP-II ENCODE K562 ChIA-PET replicate 2 data. Other replicates are shown Supplementary Figure S19. A Fisher’s exact test was computed to assess the statistical significance of the results (p-value = 1.166e-05). Similar results were observed with other CAGE and /or ChIA-PET replicates. **B.** The coordinates of the regions interacting with ‘expressed’ or ‘non-expressed’ T STR CAGEs as defined by ChIA-PET were intersected with that of the genome segments provided by combined chromHMM/Segway in K562. Fisher’s exact tests were performed to assess potential enrichments in the indicated segments (**, *<* 0.001). Similar results were observed with other CAGE and /or ChIA-PET replicates. E, ‘Predicted enhancer’; R, ‘Predicted Repressed or Low Activity region’; T, ‘Predicted transcribed region’; TSS, ‘Predicted promoter region including TSS’. **C.** The coordinates of the regions interacting with ‘expressed’ or ‘non-expressed’ T STR CAGEs were intersected with FANTOM5 CAGE coordinates and the number of CAGEs in each class was calculated. Fisher’s exact tests were performed to assess potential enrichments in the indicated classes (**, *<* 0.001).

### Transcription at T STRs can be predicted by a sequence-based deep learning model

We decided to probe transcription at T STRs using a machine learning approach. Specifically, we used a regression strategy to predict the mean raw tag count of each T STR (*>* 9 Ts) across all FANTOM libraries (Supplementary Figure S21). We first studied the link between tag count and STR length and observed that T STRs with the same length can exhibit different tag counts and, conversely, T STRs with different length can harbour similar tag count (Supplementary Figure S22A). Plotting the densities of mean raw tag counts at T STRs associated or not with CAGE peaks indicates the existence of T STRs not associated with CAGE peaks but associated with CAGE tags nonetheless (Supplementary Figure S23). We computed G bias, as in Figure 2, at T STRs not associated with CAGE peak but associated with high or low tag count (Supplementary Figure S24). As expected, T STRs without Heliscope CAGE peak and low tag count are associated with few Illumina CAGE reads compared to T STRs without CAGE peak but high tag count, despite being *>* 4 times more numerous (Supplementary Figure S24). However, Illumina reads detected at T STRs without CAGE peak but high tag count (*>* 18.45, n = 52,999) are clearly biased towards G, in contrast to T repeats without CAGE peak but low tag count (*<* 4, n = 218,074). These results support the existence of potential false negative FANTOM CAGE peaks and likely explain results depicted in Figure 2B and Supplementary Figures S9 and S11.

The machine learning approach considered was a deep Convolutional Neural Network (CNN). Model architecture and constructions of the different sets used for learning are shown in Supplementary Figure S25. As input, we used sequences spanning ± 50bp around the 3’ end of each T STR. Longer sequences were tested without improving the accuracy of the model (Supplementary Figure S26). The accuracy of our model, computed as Spearman correlation between the predicted and the observed tag counts, is remarkably high (median Spearman r over 10 trainings = 0.81, Supplementary Figure S27A) and the error low (median absolute error for one model = 1.82, Supplementary Figure S27B). Note that we also trained CNN model to predict transcription at T STRs in mouse (Spearman r = 0.77 and median absolute error = 1.82) and chicken (Spearman r = 0.61 and median absolute error = 1.09) but, as previously mentioned, compared to human, mouse and chicken CAGE data data are small-scaled in terms of number of reads mapped and diversity in CAGE libraries [2], making the signal less accurate than in human (see Supplementary Figure S28 for details).

### Probing sequence-level instructions for transcription at T STRs

We further used our deep CNN model to probe the sequence-level instructions for transcription at T STRs. We noticed that the length of the T STRs associated with CAGE peaks is greater than that of T STRs not associated with CAGE peaks but located within the same genes (Figure 6A). Likewise, the CAGE mean TPM expression, computed in [2], slightly increases with T STR length (Figure 6B). We thus used a perturbation-based approach [40] and compared CNN predictions simulating varying lengths of STRs. We chose a T STR with a low error and increased the length of its polyT track ranging from 9 to 51. The predicted tag count increases with the length of the polyT track (Figure 6C), revealing the influence of T STR length in CNN predictions. Furthermore, we randomly modified sequences in order to maximize their predicted tag count and to computationally generate an optimized sequence (see methods). Pairwise comparisons of these optimized sequences (2,000 comparisons) reveal that the main feature common to all optimized sequences is the length of the T repeat (Supplementary Figure S29).

**Figure 6.**
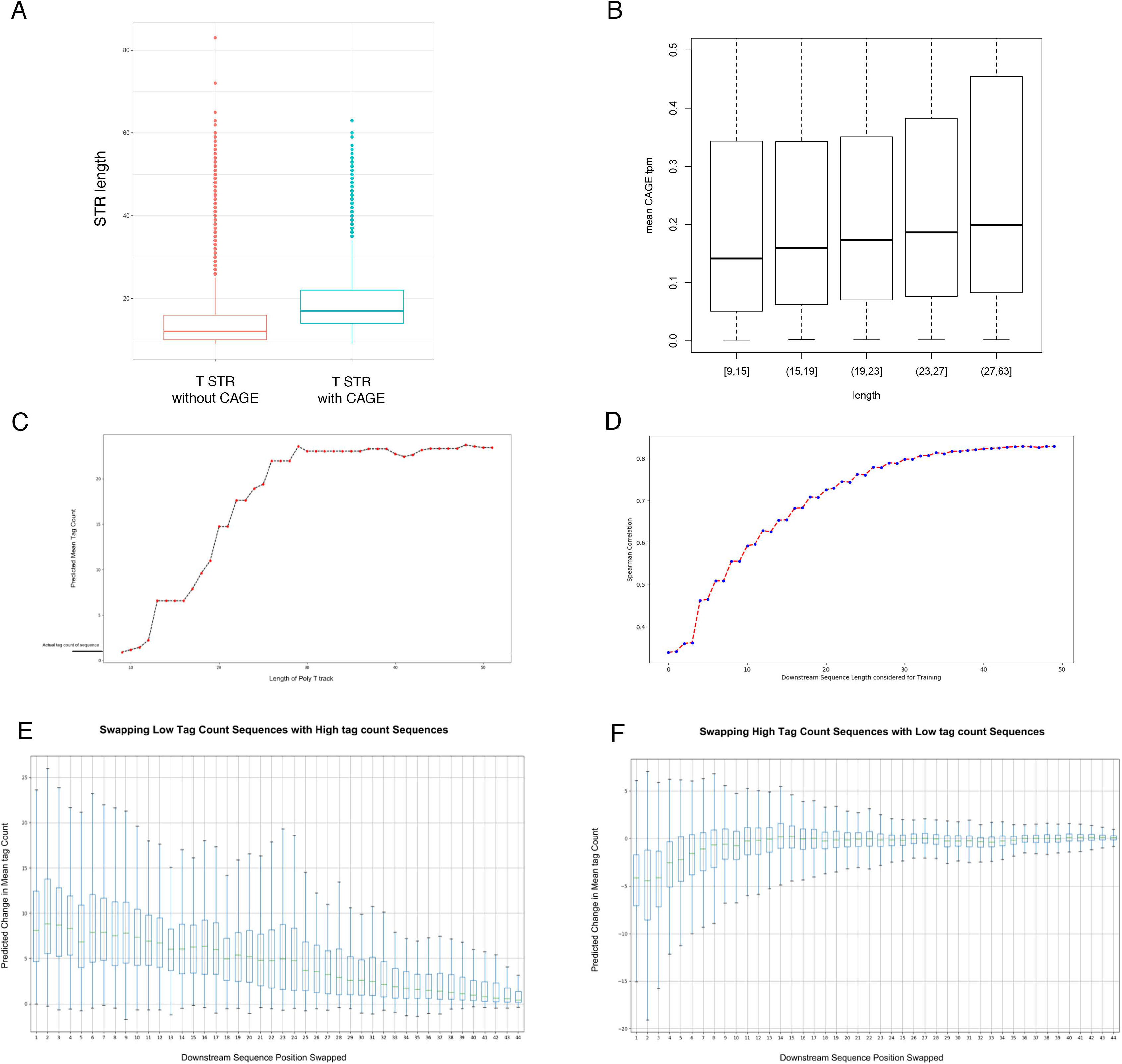
Probing sequence-level instructions for transcription at T repeats. **A.** Length of T STRs associated (blue) or not (red) with CAGE peaks. For sake of comparison, only T STRs without CAGE peak and located within the same genes as T STRs with CAGE peaks were considered. **B.** Average CAGE TPM expression computed in [2] in five T STR length intervals as defined by the R *quantcut* function. The number of CAGE peaks considered is 10,632. Y-axis was limited to 0.5. **C.** A T STR whose transcription is predicted with low error was chosen and its length was gradually increased from 9 to 51 (x-axis). The prediction was assessed at each step (y-axis). **D.** Deep CNN models were trained using 50 bp upstream T STR ends and varying lengths of sequences located downstream, ranging from 0 to 50 bp (x-axis). For each trained model, the Spearman correlation was computed on the test set (30% of entire set of T STRs i.e. 350,770 sequences) between the prediction vector and that of observed tag counts (i.e. accuracy of the model, x axis). **E and F.** 50 sequences with low error and high tag count (set ‘high’) and 50 sequences with low error and low tag count (set ‘low’) were chosen. For all pairs of ‘high’ and ‘low’ sequences, we sequentially replaced 5-mers from one sequence with 5-mer from the other and predicted the tag count of the 2 new sequences for each 5-mer. Transcription change was assessed as the difference between tag counts before and after 5-mer swapping. (E) shows the results of insertions of 5-mers of ‘high’ sequences in ‘low’ sequences, while (F) shows the results obtained inserting 5-mers of ‘low’ sequences in ‘high’ sequences.

It is important to note that the mean tag count used as input to learn our CNN model is based on CAGE data previously mapped along T STRs [2](see Supplementary Figure S30 for a representative example). Consequently, this metric may be influenced by the length of T STR itself (Supplementary Figure S22). To exclude that the importance of T STR length captured by our CNN model is merely linked to the tag count calculation, we built another model considering as input the mean tag count computed in an arbitrary window encompassing 20bp upstream and 5bp downstream T STR ends, no matter the length of the T STR. In that case, the mean tag count does not necessary reflect the CAGE signal mapped by FANTOM (see for instance Supplementary Figure S30) but it is insensitive to T STR length (Supplementary Figure S22B compared to Supplementary Figure S22A). This model yields similar accuracy (Spearman r between prediction/observation = 0.83) and the influence of T STR length is also captured, although, as expected, to a lesser extend than with the initial model (Supplementary Figure S31). Because this second model is built on an arbitrary window instead of genomic annotation (i.e. STR), and to stay in agreement with previous CAGE mapping [2], we decided to use the first model, trained on CAGE tag count computed along STR length, for the rest of the study.

We next tested whether our model learns additional features, lying downstream T STR. The accuracy of our model increases considering increasing lengths of sequence located downstream T STR with a plateau at 40-50 bp (Figure 6D). We further considered 50 sequences with low error and high tag count (set ‘high’) and 50 sequences with low error and low tag count (set ‘low’). For all pairs of ‘high’ and ‘low’ sequences, we sequentially replaced 5-mers from one sequence with 5-mers from the other. We then predicted the tag count of the 2 new sequences for each 5-mer swapping and assessed transcription change as the difference between tag counts before and after 5-mer swapping. Inserting 5-mers of ‘high’ sequences in ‘low’ sequences induces overall positive changes (i.e. increased transcription) (Figure 6E), while inserting 5-mers of ‘low’ sequences in ‘high’ sequences overall decreases transcription (Figure 6F). Besides, these 5-mer segmentation reveals that the instructions located near the end of T STRs (i.e. close to the CAGE peak summit) are crucial (i.e. can induce more change in the prediction) compared to the rest of the sequence (Figures 6E and F). Looking for motif enrichment, we further noticed that several motifs are enriched in T STR with high tag counts (top 5,000) compared to low tag counts (bottom 5,000) (Supplementary Figure S32). However, except the T homopolymer (Supplementary Figure S32), no single motif clearly distinguishes T STRs with high tag count from T STRs with low tag count. Likewise, a simple CNN with only one convolutional layer (1 layer-CNN), which is similar to finding enriched k-mers or motifs, poorly performs (median Spearman r = 0.2). Hence, a single representation cannot model the intricate structure of the features captured by CNN to predict transcription at T STRs.

Alternative methods were also tested, namely LASSO [41] and Random Forest [42], using, as predictive variables, mono-, di- and tri-nucleotides rates in the region located upstream (−50bp) or downstream (+50bp) T STR ends, the T STR lengths and the first nucleotide (A,C,G or T) downstream the T STR (−1 when considering 0 as the CAGE summit). In that case, the model accuracies are 51% and 61% for LASSO and Random Forest respectively. These models are not as accurate as CNN but they confirm that T STR length is the main feature being used from the sequence located upstream T STR end, while other features lie downstream the T STR (Supplementary Table S4 and Supplementary Figure S33). These analyses confirm the existence of instructions for transcription lying downstream T STRs, arguing against artifactual internal priming.

### Clinically relevant variants are predicted to impact transcription at T STRs

Previous studies showed that genomic variations at T STRs can impact gene expression. For instance, the variant rs10524523, located in TOMM40 intron 6, which corresponds to a T STR associated with a CAGE signal in several FANTOM libraries, is linked to the age of onset of cognitive decline, Alzheimer’s disease and sporadic inclusion body myositis [43–48]. The TOMM40 mRNA brain expression appears to be linked to the length of the T STR, with the longer the variant the higher the expression [43, 44]. Conversely, the KIAA0800/ENSG00000145041 promoter is repressed by an upstream element involving a T STR and two Alu repeats [49]. In that case, the CAGE signal detected at this particular T STR has been annotated as one TSS of KIAA0800/ENSG00000145041 in FANTOM CAT (namely ENCT00000303320.1, FTMT20900010597.1, MICT00000243080.1) [4]. More widely, the length of T STRs have been reported to impact gene expression (called expression STR or eSTR) in a directional and strand-specific manner [50].

We further used our model to predict the effects of genomic variations located within sequences surrounding T STR (end of T STR ± 50bp). First, we observed, in the case of TOMM40 intron 6 variant, that predicted transcription is positively correlated to the length of the T STR (Figure 7A). Second, we compared the effect of variants listed in dbSNP, the world’s largest database for nucleotide variations [51], and variants listed in ClinVar, which pinpoints medically important variants [52]. Almost all T STRs harbor variations listed in dbSNP (962,241 out of 1,169,145), while 2,577 T STRs are associated with variations listed in ClinVar. Strikingly, these T STRs tend to be associated with high tag counts compared to T STRs with dbSNP variants (Figure 7B), indicative of potential clinical relevance of transcription at T STRs. The ClinVar variants were not equally distributed around T STRs (Figure 7C). Overall, ClinVar variants are frequently found around T STRs, with a peak at position 0 and +1 (0 being the end of T STR), close to the CAGE summit (+2 in that case). This particular distribution was not observed when considering all permissive CAGE peak summits (Supplementary Figure S34). The variants located at the vicinity of T STRs are predicted by our model to induce extensive changes in transcription, being either positive (i.e. transcription increase) or negative (i.e. transcription decrease) when located at the vicinity of T STR, and overall negative when located within T STR (Figure 7D). Genomic variations within a T STR will more likely disrupt the repeat and decrease its length, thereby inducing an overall decrease in transcription prediction. The predicted changes are not linked to the nature of the variants (i.e. SNP or insertion/deletion) as any types of variants can have positive and negative effects on transcription with insertions/deletions having broader effects (Supplemental Figure S35). Variations linked to certain diseases were enriched in the 101bp-long regions centered around T STR ends, but no clear enrichment for a specific type of diseases was noticed (Supplementary Table S5).

**Figure 7.**
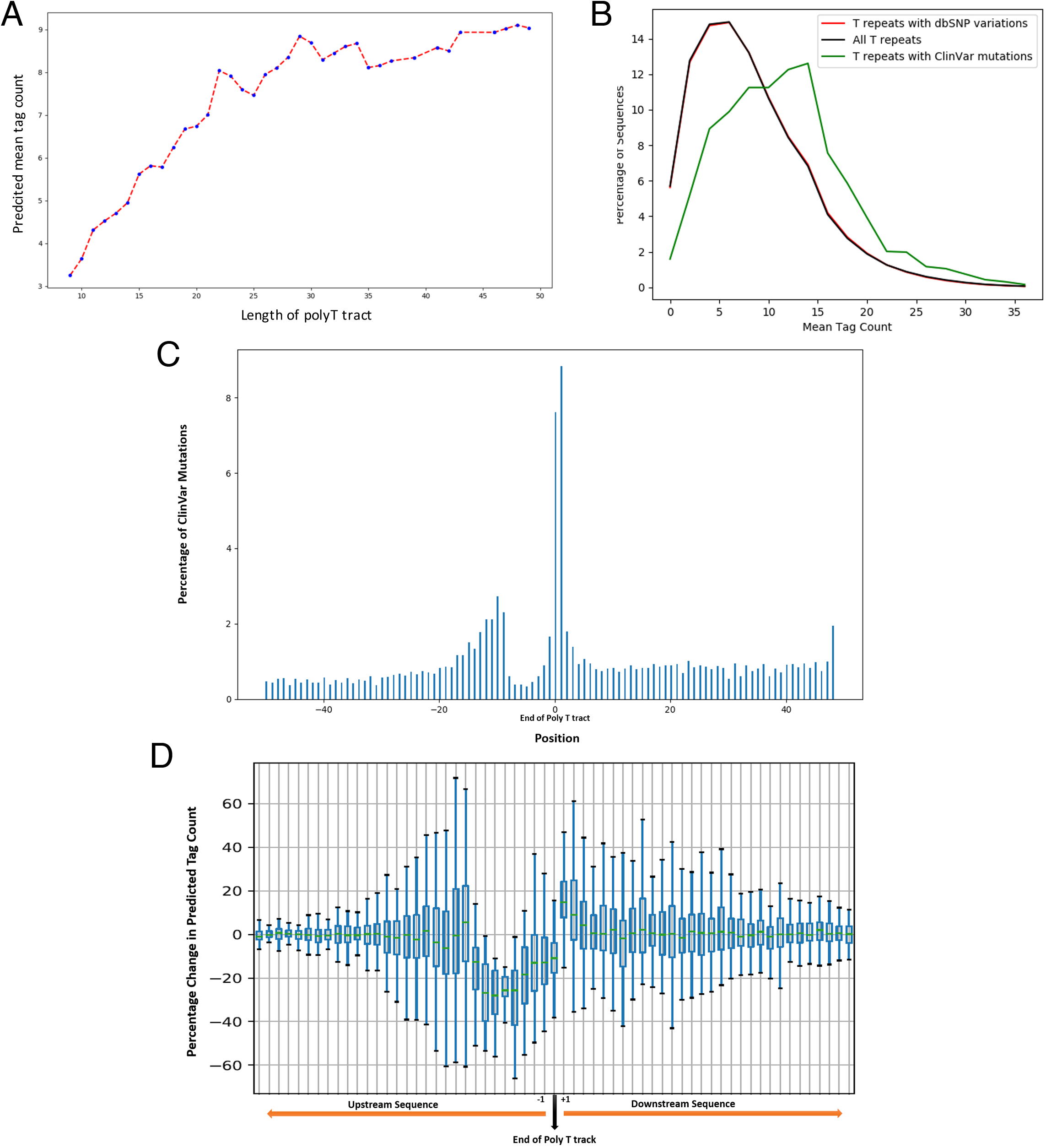
Evaluating the effect of genetic variants on transcription at T STRs. **A.** Increasing the length of the T STR located in TOMM40 intron 6 as in Figure 6C increases the predicted tagcount. **B.** Tag count distribution of all T STRs (black), T STRs with dbSNP variants (red) and T STRs associated with ClinVar variants (green). **C.** Frequency of ClinVar variants around T STR ends (position 0). The y-axis shows the fraction of all ClinVar variants located at the position indicated on the x-axis. Low variant frequency within T STRs is presumably due to mapping issues. **D.** Effect of prediction changes induced by ClinVar variants (see methods for details). The changes are shown as percentages of the original tag count in a position-wise manner (see ‘methods’ section).

## Discussion

We looked at recurring DNA motifs associated with unassigned FANTOM CAGE peaks [2] and showed that a significant portion of them initiate at T STRs in several species. We do not exclude that some CAGE peaks initiating at T STRs are generated by internal priming during Heliscope sequencing. Yet we provide evidence that a fraction represents genuine TSSs as (i) T STR CAGEs are truly capped and associated with typical epigenetic marks, (ii) several of them correspond to annotated long non-coding transcripts or enhancer RNAs and are confirmed by various technologies, not requiring oligodT priming, and (iii) their expression can be predicted by a deep learning model, which uses features located downstream T repeats, unlikely involved in internal priming. We show that GRO-seq coincides with CAGE at T STRs (Figure 3B and Supplementary Figures S11 and S12). Li *et al.* suggested that CAGEs detected at polyA/T repeats correspond to technical artifacts because they are not detected by a GRO-seq method that enriches for 5’-capped RNAs (GRO-cap) [22]. However, in comparison to CAGE, GRO-cap generates fewer reads mapping to introns [53], making hard to confirm the existence of faintly expressed and intronic CAGEs, such as T STR CAGEs, with this technology. In fact, non-coding transcription appear overall much less efficient than that of mRNAs [7] (Supplementary Figure 7A). It is also worth noticing that several features insinuating that T STR CAGEs are technical artifacts may constitute specific properties of non-coding RNAs with, notably, the majority of lncRNAs being enriched in the nucleus [54–56].

The human genome is scattered with repetitive sequences, and the vast majority of the genomic DNA is supposed to be transcribed [1, 5, 6]. Implicitly, the human transcriptome should contain a large portion of RNAs derived from repetitive elements [57]. We provide here an example of such RNAs, which may represent the counterparts of satellite non-coding RNAs [58].. Homopolymers of *>* 10 Ts appear at frequencies well above chance in the non-coding regions of several organisms, suggesting selective advantages that lead to their overrepresentation [26]. Additionally, using FANTOM and GENCODE genome annotations, we noticed that T STRs are enriched in introns of coding, as opposed to non-coding, genes (Supplementary Figures S1 and S2).

In human and mouse genomes, T STRs can correspond to the polyA tails of short interspersed nuclear elements (SINEs) [59]. According to RepeatMasker, only a fraction of human T STRs associated with CAGE peaks correspond to SINEs (∼ 37% vs. ∼ 50% for T repeats without CAGE peak, Supplementary Table S6). T STR CAGEs are also detected in chicken (Supplementary Figure S1) where SINE elements are rare [60] (note that repeats of ≥ 9 Ts are as abundant in chicken and human and represent 0.04% of the genome). These observations support the idea that transcription at T STRs exists independently of SINE elements. However, we do not exclude a possible co-evolution between T STR transcription and SINEs, at least in human and mouse, notably because of the importance of polyA tail length in SINE transposition in one hand [61, 62] and the influence of this length on transcription revealed in this study in the other hand.

At this stage, whether T STR-associated non-coding RNAs or the act of transcription *per se* is functionally important [7] remains to be clarified. This is all the more warranted as T STR transcription is associated with clinically relevant genomic variations (Figure 7B). This non-coding transcription, mostly sense and intronic, may specifically regulate certain coding - as opposed to non-coding - genes (Supplementary Figure S7, Supplementary Tables S1 and S2). In line with this idea, we show here that T STR CAGEs preferentially interact with gene TSSs (Figure 5).

T homopolymers have been shown to act as promoter elements favoring transcription by depleting repressive nucleosomes [63], specifically by orientating the displacement of nucleosomes [64]. As a consequence, T homopolymers can increase transcription of reporter genes in similar levels to TF binding sites [65]. Besides, eSTRs located within the gene body are more likely to have T on the template strand and show higher gene expression with increasing repeat number [50]. This directional and strand-specific bias [50], which cannot be explained by mere nucleosome depletion [63, 64], could very well be molecularly caused by transcription. These observations, in addition to overrepresentation of long T STRs in eukaryotic genomes [26], suggest functional role(s) for transcription at T STRs. Dedicated experiments are now required to formally assess its overall contribution in gene expression.

## Methods

### Datasets and online resources

Publicly available data used in this study can be found at the urls provided in Supplementary Table S7. The genome assemblies considered are human hg19, mouse mm9, chicken galGal5, dog canFam3, rat rn6, rhesus macaque rheMac8. Human Introns and RepeatMasker coordinates were downloaded from UCSC table browser.

### Motif analyses

The HOMER motif analysis tool [23] was used to find 21bp long-motifs in regions spanning 10bp around CAGE peakmax (*findMotifsGenome.pl* with options -len 21, -size given, -noknown and -norevopp). HOMER was further used to identify, CAGEs harboring specific motifs (*annotatePeaks.pl* with option -m) in particular T homopolymers (motif#2 in Figure 1).

### Characterization of T repeats in human and mouse

T STRs of *>* 9Ts were identified in the human, mouse and chicken genome using the tool developed by Martin *et al.* [66]. The human, mouse and chicken coordinates are provided as bed files available at https://gite.lirmm.fr/ibc/t-repeats.

### Bioinformatics tools

Intersections between genomic coordinates were computed with BEDTools [67]. Signal were extracted from bigwig files using bwtool (*agg* and *extract* subcommands) [68] and deepTools [69]. Statistical tests were done using R [70] (*fisher.test, wilcox.test, cor*) or Python (*fisher exact, mannwhitneyu, spearmanr* from scipy.stats (http://www.scipy.org/)). Graphs were made using ggplot2 R package [71] or Python matpotlib library (https://matplotlib.org/api/pyplotapi.html). All scripts are available upon request.

### Evaluating mismatched G bias at Illumina 5’end CAGE reads

Comparison between Heliscope vs. Illumina CAGE sequencing was performed as in de Rie *et al.* [31]. Briefly, ENCODE CAGE data were downloaded as bam file (Supplementary Table S7) and converted into bed file using samtools view [72] and unix awk as follow:

~~~
**samtools view file.bam | awk ‘{FS=“\t”}BEGIN{OFS=“\t”}{if($2==“0”) print $3,$4-1,$4**,
   $10,$13,“+”; else if($2==“16”) print $3,$4-1,$4,$10,$13,“-”}’ > file.bed
~~~

The bedtools intersect [67] was further used to identify all CAGE reads mapped at a given position (see Figure 2B and C). The unix awk command was used to count the number and type of mismatches as follow:

~~~
intersectBed -a positions_of_interest.bed -b file.bed -wa -wb -s |
awk ‘{if(substr($11,1,6)=="MD:Z:0" && $6=="+") print substr($10,1,1)}’ | grep -c "N"
~~~

with N = {A, C, G or T}, positions of interest.bed being for instance T STR CAGE coordinates and file.bed being the Illumina CAGE tag coordinates.

Absence of mismatch focusing on the plus strand were counted as:

~~~
intersectBed -a positions_of_interest.bed -b file.bed -wa -wb -s |
awk ‘{if(substr($11,1,6)!="MD:Z:0" && $6=="+") print $0}’ | wc -l
~~~

As a control, 61,907 random positions were generated with bedtools *random* (−l 1 and -n 61907). We also used the 3’ end of the pre-miRNAs, which were defined, as in de Rie *et al.* [31], as the 3’ nucleotide of the mature miRNA on the 3’ arm of the pre-miRNA (miRBase V21, see Supplementary Table S7), the expected Drosha cleavage site being immediately downstream of this nucleotide (pre-miR end + 1 base).

### Epigenetic marks at T repeats

Average chromatin CAGE signal around T STR CAGEs (Figure S9) were computed using the *agg* subcommand of the bwtool [68]. For RNAP-II ChIA-PET, interacting regions within the same chromosome were extracted using the column name $4 of the bed file. Roadmap and GENCODE ChIP-seq as well as GROseq data were extracted from URLs indicated in Supplementary Table S7. To look at epigenetic marks around T repeats into H1-hESC cell line, the coordinates of ‘expressed’ and ‘non-expressed’ T repeats ends+2 bp (see results for details) were intersected with Roadmap bedfiles using bedtools intesect [67].

~~~
intersectBed -wb -a epigenetic_mark.bed -b expressed_polyT_CAGEs.bed | sort -u | wc -l
~~~

The fractions of intersecting T repeats within each set were compared for each available epigenetic mark and each of the 3 H1-hESC samples using Fisher’s exact test. The same procedure was used in other cell lines with ENCODE broadpeaks data for the H3K4me1, H3K4me2, H3K4me3 and H3K36me3. For GRO-seq data, we used bigwig files from the ‘gro-seq.colorado’ repository (Supplementary Table S7). In H1-hESC cell line, we compared the fraction of intersecting ‘expressed’ and ‘non-expressed’ T repeats similar to the procedure used for GENCODE and Roadmap data. The GRO-seq signal around these two sets was also compared as the mean signal into a 100 bp window centered on the T repeat end using deeptools multi-BigwigSummary [69].

~~~
multiBigwigSummary BED-file -b CellLine_Nascent_RNAseq_files_forward/*.bw
  -o out.npz --BED sample_polyT.end_minus.plus50_coordinates_plusStrand.bed --
    outRawCounts sample_polyT.end_minus.plus50_meanSignal_plusStrand.txt
~~~

~~~
multiBigwigSummary BED-file -b CellLine_Nascent_RNAseq_files_backward/*.bw
  -o out.npz --BED sample_polyT.end_minus.plus50_coordinates_minusStrand.bed --
    outRawCounts sample_polyT.end_minus.plus50_meanSignal_minusStrand.txt
~~~

We compared the fractions of T repeats with a no-null signal into both ‘expressed’ and ‘non-expressed’ sets using Wilcoxon test. We also calculated the sum of these mean signals across all the samples for each T repeat with similar results.

### Convolutional Neural Network

A Zenbu [73] xml custom script was used to calculate the mean raw tag count (Q3 filter, no normalization) of each base genome-wide in 988 FANTOM libraries. We summed these means along each T STR + 5bp. This values were used as predicted variables in a CNN model built on sequences centered around each T repeat end (n = 1,169,236). The same procedure was repeated with mouse (397 libraries) and chicken data (58 libraries) using the same architecture. The coordinates of the T repeats with the mean tag count values (score column) used to train our models are provided as bed files https://gite.lirmm.fr/ibc/t-repeats. The fasta file corresponding to T repeats sequences with their tag count was parsed using Biopython [74] library. To build the convolutional neural network, we used Keras [75] library in Python with Tensorflow [76] backend. The subsequent analyses of altering length of T repeats and swapping downstream sequence was performed by treating sequences as strings. The complete code can be found at https://gite.lirmm.fr/ibc/t-repeats.

### Feature extraction

We looked at motif by random optimization of the CNN prediction using the following algorithm: first, two sequences are randomly chosen in the test set. For each sequence, the effect of changing one nucleotide at a random position into A,T, C or G is assessed. The nucleotide with the maximum predicted tag count is kept and the procedure continues testing another position. This procedure is repeated until no further increase of predicted tag count is observed over 3,500 randomly selected positions. The two optimized sequences are then compared position-wise in order to return 1 for each position with the same nucleotide in both sequences and 0 otherwise. This process was repeated for 2,000 pairs of random sequences. Same procedure was applied minimizing the tag count for each sequence. The sequence logos were generated using the *seqlogo* library https://pypi.org/project/seqlogo/. The complete code can be found at https://gite.lirmm.fr/ibc/t-repeats.

### Predicting impact of ClinVar variants

ClinVar and dbSNP vcf files were downloaded and then converted into bed files.They were intersected with coordinates of all T repeats using bedtools intersect [67] as follows:

~~~
bedtools intersect -a clinvar_mutation.bed -b t_repeats.bed -wa -wb >
    t_repeats_clinvar.bed
bedtools intersect -a db_snp_mutation.bed -b t_repeats.bed -wa -wb > t_repeats_dbSNP
    .bed
~~~

The bed files generated above were then converted into Pandas [77] dataframe in python and variants were introduced in T repeats sequences. The CNN model developed previously was then used to predict the mean tag count of the mutated sequences. The change was computed using the following formula:

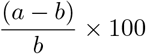

where *a* is the predicted tag count after variation and *b* the predicted tag count before variation. The complete code can be found at https://gite.lirmm.fr/ibc/t-repeats.

### Comparison with other models

To compare our deep learning regression model with other parametric and non parametric approaches, we used linear regression with l1-norm penalty via LASSO [41] and Random Forest [42] to predict the mean tag count signal. For both approaches, the predictive variables are the nucleotide, dinucleotide and trinucleotide content computed into the 50 bp downstream and 50 bp upstream the T homopolymer end, excluding the T homopolymers. We also integrated the length of the T repeat and the nature of the nucleotide following the T repeat end (A, C or G at position −1 referring to the CAGE summit at 0). LASSO inference was performed using the function cv.glmnet from the R package glmnet and Random Forest was performed using the randomForest R package. To compute the distribution of the minimal depth we used the *min depth distribution* function from the R package randomForestExplainer.

## Declarations

### Ethics approval and consent to participate

Not applicable.

### Consent to publish

Not applicable.

### Availability of data and materials

All data used in this study are provided in Supplementary Tables. The code to predict transcription at T STRs with a CNN is available at https://gite.lirmm.fr/ibc/t-repeats.

### Author’s contributions

C.B., M.S., M.G. W.W.W., M.d.H, L.B and C-H.L. analyzed and interpreted data. M.S. and M.G. developed CNN models and extracted features. C.M. identified T repeats in human, mouse, chicken genome. J.R., Y.H., A.H., H.S., S.N., I.M. generated CAGE data used in this study. M.d.H., J.S. and C-H.L. generated Zenbu tracks. M.d.H and C-H.L. studied G bias at ENCODE read 5’ ends. P.C., W.W.W, L.B. and C-H.L acquired fundings. C-H.L. wrote the manuscript. All authors have read and approved the manuscript.

## Acknowledgements

We thank Cédric Notredame, Anthony Mathelier, Oriol Fornes Crespo, Philip Richmond, Jean-Christophe Andrau, Diego Garrido Martin, Dimitri D. Pervouchine, Roderic Guigo, Charles Plessy and Chung Hon for their help in analyzing the data and for insightful suggestions. We are indebted to the researchers around the globe who generated experimental data and made them freely available. C-H.L. is grateful to Marc Piechaczyk and Edouard Bertrand for continued support. The work was supported by funding from CNRS (International Associated Laboratory ”miREGEN”), INSERM-ITMO Cancer project ”LIONS” BIO2015-04, *Plan d’Investissement d’Avenir* #ANR-11-BINF-0002 *Institut de Biologie Computationnelle* (young investigator grant to C-H.L.) and GEM Flagship project funded from Labex NUMEV (ANR-10-LABX-0020). FANTOM5 was made possible by the following grants: Research Grant for RIKEN Omics Science Center from MEXT to Y.H.; Grant of the Innovative Cell Biology by Innovative Technology (Cell Innovation Program) from the MEXT to Y.H.; Research Grant from MEXT to the RIKEN Center for Life Science Technologies; Research Grant to RIKEN Preventive Medicine and Diagnosis Innovation Program from MEXT to Y.H. This work was further supported by a Research Grant from MEXT to the RIKEN Center for Integrative Medical Sciences.

## Additional Files

Additional figures

**Figure S1 Motif discovery around CAGE summit.** The procedure described in Figure 1 was repeated with CAGE peak summits collected in chicken, dog, macaque and rat.

**Figure S2 Heliscope vs. Illumina CAGE sequencing. A.** In the HeliScopeCAGE protocol, capped RNA will be randomly primed and the RNA/DNA duplex will be captured on beads by the CAP Trapper reaction. The first-strand cDNAs are then released and a poly-A tail is added to prepare them for sequencing. The HeliScope platform primes the sequencing reactions with poly-dT oligonucleotides grafted to the surface of its flow cells. In the HeliScope CAGE, it is intended that the sequencing reactions will start after the poly-A tail added to the first-strand cDNA, thus producing reads that align at the TSS. If the first-strand cDNA contains internal A-rich regions, the priming can also happen internally and yield CAGE peaks at internal poly-T tracts. **B.** Comparison of FANTOM (Heliscope) and ENCODE (Illumina) CAGE peaks. Four examples are shown. Top, FANTOM ; middle, ENCODE ; bottom, FANTOM CAGE peaks; green: (+) strand; purple: (−) strand.

**Figure S3 Motif discovery around Start-seq and DECAP-seq peak summits. A.** The procedure described in Figure 1 was repeated on TSSs detected by Start-seq [27] (forward data, n = 1,086,787) normalizing by total GC-content with -gc option. B. Same procedure applied on DECAP-seq TSSs [28] (n = 106,742). GSM2422532 and GSM2422533, which are not stranded data, were merged. In that case, the INR motif is not detected with this procedure and motif 4 (poly tract) is labelled as possible false positive. Note that the number of mouse CAGE summits [2] located within a 10bp window around Start-seq [27] and DECAP-seq [28] TSSs is low: ~ 34.7% of CAGE TSSs have same coordinates as Start-seq TSSs but only ~ 6.8% of CAGE TSSs share coordinates with DECAP-seq TSSs and ~ 7.2% of Start-seq TSSs share coordinates with DECAP-seq TSSs. However, we noticed that mouse STRs of ≥ 9Ts associated with CAGE peaks are more associated with Start-seq and DECAP-seq peaks than T STRs not associated with CAGE peaks: ∼ 2% of CAGE-associated T STRs are also associated with Start-seq peaks (94 out of 4,600 T STRs considering only forward data) with only ∼ 0.3% T STRs not associated with CAGE but associated with Start-seq peaks (1256 out 413475, Fisher’ exact test p-value < 2.2e-16). Likewise, ~ 0.36% of CAGE-associated T STRs are also associated with DECAP-seq peaks (32 out 8825) with only ~ 0.03% T STRs not associated with CAGE but associated with DECAP-seq peaks (307 out 826129, Fisher’s exact test p-value < 2.2e-16).

**Figure S4 Motif discovery around TSS-seq peak summits collected in Arabidopsis thaliana.** The procedure described in Figure 1 was repeated on TSS-seq peaks [29] normalizing by total GC-content with -gc option.

**Figure S5 Distribution of CAGE peaks at homopolymers of A/T A.** 81,080 STRs from STR reference catalog [14] are associated with a CAGE peak (considering a window of 5bp around the STR): 36,062 correspond to homopolymers of Ts and 33,802 to homopolymers of As. Most CAGE peaks associated with STRs of T are on the ‘+’ strand (35,653 out of 36,185), while CAGE peaks associated with homopolymers of As are on the ‘-’ strand (33,313 out of 33,922) B. CAGE peaks are not equally distributed around polyA/T STRs with 35,974 and 686 located respectively downstream and upstream of polyT STRs (CAGE tags within STRs are here counted in both cases explaining why 35,974+686 > 36062) and 679 and 33,723 located respectively downstream and upstream of polyA STRs. Hence, CAGE peaks are almost exclusively located downstream polyT STRs. Since no amplification is involved in CAGE sequencing [20], imbalance for polyT vs. polyA supports the idea of internal priming (note that FANTOM CAT genes contain 628,762 T repeats and 499,419 A repeats).

**Figure S6 T STR CAGEs harbor specific features. A.** Directionality, extracted from [4], of all CAGEs (red) and that of 10,926 T STR CAGEs (blue) are shown as boxplots. Directionality close to 1 indicates that sense transcription is more abundant than antisense transcription within a region of −800bp/+200bp centered around the CAGE summit while directionality of −1 indicates that antisense transcription is more abundant. These results argue against the presence of divergent (i.e. upstream antisense) RNAP-II transcription associated with T STR CAGEs, as widely observed for canonical TSSs [78] **B.** Exosome sensivity score were extracted from [4]. Exosome sensitivity of a CAGE cluster is measured as the relative fraction of CAGE signal observed after exosome knockdown in HeLa-S3 cells as previously described in [30]. The exosome sensitivity of all CAGEs (red) and that of 10,926 T STR CAGEs are shown as boxplots.

**Figure S7 T STR CAGEs are faintly expressed and detected in highly expressed genes. A.** Expression medians were computed for each permissive CAGE, CAGE assigned to genes and T STR CAGEs across 1,829 libraries. The Phase 1 and 2 combined Tag Per Million (TPM) normalized data were used (Supplementary Table S7). The y-axis was limited to 20. Overall T STR CAGEs appear faintly expressed (median of median CAGE TPM expression in all samples = 0.3347) compared to all CAGEs detected (median = 1.117, Wilcoxon test p-value < 2.2e-16) or to CAGEs assigned to gene TSS (median = 2.396, Wilcoxon test < 2.2e-16) **B.** T STR CAGEs are preferentially detected in highly expressed genes. Distribution of median expression was computed across 1,829 FANTOM5 libraries of genes containing T repeats with (blue) or without (purple) CAGE (Wilcoxon test p-value < 2.2e-16). The median expressions of all FANTOM CAT genes (green) and that of PCGs (red) are shown for sake of comparison. The y-axis was limited to 40.

**Figure S8 T STR CAGEs are preferentially detected in the nuclear compartment.** For each indicated library, CAGE expression (RLE normalized) was measured in nuclear and cytoplasmic fractions. Each CAGE was then assigned to nucleus, cytoplasm or both compartments. The number of CAGEs in each class is shown for each sample as a fraction of all detected CAGEs. The sample Fibroblast Skin 2 likely represents an outlier. **A.** 63,974 CAGEs associated with T repeats **B.** 1,048,124 CAGEs detected **C.** 130,286 CAGEs assigned to PCGs **D.** 68,731 CAGEs harboring the third motif shown in Figure 1.

**Figure S9 T STR CAGEs are associated with chromatin.** Distribution of chromatin-associated RNAs extracted from K562 cells around T STRs with (green) or without (blue) CAGE peaks. Total CAGE signal on + strand is shown. Data are from ENCODE and were generated using Illumina technology. Chromatin-associated RNAs are detected even at T STRs without CAGE peaks, suggesting the existence of false negatives (see Supplementary Figure S24).

**Figure S10 Motif discovery around intragenic and intergenic CAGE summit.** HOMER [23] was used to find 21bp-long motifs in 21bp-long sequences centered around the summit of FANTOM5 CAGEs located (intragenic) or not (intergenic) in FANTOM CAT genes. Intergenic CAGE summits were used as foreground, while intragenic summits were used as background. The motif shown is found in only 0.13% of the intergenic CAGE summits, making it hardly discriminative.

**Figure S11 G bias at T STR CAGEs.** G bias in ENCODE CAGE reads was assessed at −2 of CAGE peakmax as in Figure 2 in GM12878 (A), K562 (B) and A549 (C and D).

**Figure S12 Intersection with GRO-seq peaks.** Intersection of ‘expressed’ (CAGE+)/’non expressed’ (CAGE-) T STR CAGEs (end ± 50bp) with nascent RNAseq peaks from 3 ‘gro-seq colorado’ samples (SRR*) obtained in H1 embryonic stem cells (CNhs13964, CNhs14067 and colorado’ samples (SRR*) obtained in H1 embryonic stem cells (CNhs13964, CNhs14067 and CNhs14068 FANTOM libraries). Fisher’s exact tests indicate that the fraction of T STR CAGEs with no-null GROseq signal obtained for ‘expressed’ T STR CAGEs is invariably higher than that obtained for ‘non expressed’ T STR CAGEs (p-value < 2.2e-16).

**Figure S13 Intersection with GRO-seq signal.** Intersection of ‘expressed’ (CAGE+)/’non expressed’ (CAGE-) T STR CAGEs (end ± 50bp) with nascent RNAseq peaks from 3 ‘gro-seq colorado’ samples (SRR*) obtained in H1 embryonic stem cells (CNhs13964, CNhs14067 and colorado’ samples (SRR*) obtained in H1 embryonic stem cells (CNhs13964, CNhs14067 and CNhs14068 FANTOM libraries). The sum of the mean signal for each of the ‘expressed’ (CAGE+) and ‘non expressed’ (CAGE-) T STR CAGEs is represented on the y-axis. For instance, in CNhs13964, signal was computed for 2,013 ‘expressed’ T STR CAGEs and 2,001 ‘non expressed’. Wilcoxon test was used to asses significant difference (p-value is indicated).

**Figure S14 Intersection with Roadmap epigenetics ChIP-seq data.** Fraction of ‘expressed’ / ‘non expressed’ T STR CAGEs in H1-hESC cells intersecting with H3K4me1 (top), H3K4me2 (middle), H3K4me3 (middle) and H3K36me3 (bottom) Roadmap ChIP-seq data.

**Figure S15 Intersection with ENCODE epigenetics ChIP-seq data.** A. Fraction of ‘expressed’/’non expressed’ T STR CAGEs within each library considered. B-E. Fraction of ‘expressed’ / ‘non expressed’ T STR CAGEs intersecting with H3K4me1, H3K4me3 and H3K36me3 ENCODE ChIP-seq data in H1-hESC (B), Hela-S3 (C), GM12878 (D) and K562 (E). The 3 replicates of each cell line were considered in each case.

**Figure S16 Usage and contribution of polyT-associated FANTOM CAT TSSs.** We first assessed the position of polyT-associated TSSs among each promoter: 937 TSSs out of 10,606 correspond to the most upstream TSS, arguing against internal priming and artifact hypothesis in these cases. **A.** We evaluate the usage of FANTOM CAT TSSs associated (red) or not (blue) with poly tract for each gene with more than 1 TSS. For each assigned TSS, we counted the number of samples wherein the considered TSS is expressed. We then sorted all TSSs for each gene according to this usage value and divided the rank of each polyT-associated TSS by the length of the list of all TSS assigned to each gene (usage ratio indicated in y-axis). TSSs associated with polyT repeats are slightly less used than TSSs not associated with polyT repeat (Wilcoxon test p-value < 2.2e-16). However, the median usage ratio for polyT-associated TSSs is 0.5, indicating that these TSSs are far from being the least used TSSs. **B.** We further evaluated the contribution of polyT-associated TSSs in gene expression. For each gene in each library, we divided the sum tag count (CPM RLE normalized) of all TSS associated (red) or not (blue) with polyT tract by the sum tag counts of all TSSs assigned to the gene considered (contribution ratio indicated in y-axis). As expected provided faint expression of T STR CAGEs, polyT-associated FANTOM CAT TSSs contribute only poorly to gene expression with a median contribution ratio of 0 (the median contribution of TSSs not associated with polyT repeat logically equals 1).

**Figure S17 No specific Open Reading Frame could be detected 2kb downstream polyT CAGEs.** Stop codon distribution was assessed along 2,000 nt downstream polyT-associated CAGE summits (n = 63,974). The x-axis represents the distance to transcript start and the y-axis represents the frequency of the three stop codons TAA, TAG and TGA. All possible 6 frames were considered (see y-axis labels).

**Figure S18 Motif discovery around FANTOM mouse enhancer TSSs.** HOMER [23] was used to find 21bp-long motifs in 21bp-long sequences centered around TSSs of 44,459 mouse enhancers defined in [3] and corresponding to 88,918 TSSs. Only the top 5 motifs are shown.

**Figure S19 Regions interacting with T STR CAGEs**. The coordinates of the regions interacting with ‘expressed’ or ‘non-expressed’ T STR CAGEs in CNhs12335 (A) and CNhs12336 (B) were intersected with that of the genome segments provided by combined chromHMM/Segway in K562. Fisher’s exact tests were performed to assess potential enrichments in the indicated segments (**, *<* 0.001). E, ‘Predicted enhancer’; R, ‘Predicted Repressed or Low Activity region’; T, ‘Predicted transcribed region’; TSS, ‘Predicted promoter region including TSS’.

**Figure S20 TSSs interacting with T STR CAGEs**. The coordinates of the regions interacting with ‘expressed’ or ‘non-expressed’ T STR CAGEs in CNhs12335 (A) and CNhs12336 (B) were FANTOM5 CAGE coordinates and the number of CAGEs in each class was calculated. Fisher’s exact tests were performed to assess potential enrichments in the indicated classes (**, *<* 0.001; *, *<* 0.05).

**Figure S21 Example of signal used to train CNN.** To compute the raw mean tag count of each T repeat, we first calculated the mean tag count of each base genome-wide in 988 FANTOM libraries (upper track). We then summed these values along the T repeat + 5bp (bottom track).

**Figure S22 Distribution of tag counts per T repeat length.** The raw mean tag counts are shown as boxplots according to T repeat length. The length is indicated on the x-axis. The number in bracket indicates the number of T repeats considered. Only length class of *>* 100 T repeats are shown. **A.** The tag count is computed in a window encompassing T repeat length + 5bp. **B.** The tag count computed in an arbitrary window encompassing 20bp upstream and 5 bp downstream T repeat end. This tag count computation does not consider the whole T repeat sequences but is not influenced by it lengths.

**Figure S23 Distribution of mean tag count along T repeats.** To compute the raw mean tag count of each T repeat, we first calculated the mean tag count of each base genome-wide in 988 FANTOM libraries. We then summed these values along the T repeat + 5bp. The distribution of these tag counts are plotted for T repeats initially defined as associated (red) or not (blue) with CAGE peaks as defined in [2]. The overlapping area likely represents the existence of potential false negatives i.e. T repeats not associated with CAGE peaks but associated with CAGE tags nonetheless. The y-axis was limited to 100 for sake of clarity.

**Figure S24 G bias at T STRs without FANTOM CAGE peak.** G bias in ENCODE CAGE reads (nuclear fraction, polyA-) was assessed at −2 of CAGE peakmax as in Figure 2 in A549 (two replicates), GM12878, HelaS3 and K562. T STRs not associated with CAGE peak but associated with high (tag count > 18.45, n = 52,999) or low tag (tag count < 4, n = 218,074) count were distinguished. These thresholds were defined as the median of tag count observed in T STRs with CAGE (18.45) and the first quartile of tag count observed in the case of T STRs without CAGE (4).

**Figure S25 Deep learning to predict transcription at T STRs. A.** CNN model architecture. The number and size of the filters used for convolution is indicated (’50*5’ indicates 50 filters of size 5bp; ‘30*3’ indicates 30 filters of size 3bp). B. Constructions of the different sets for learning. Only T STRs with at least 1 tag and no undetermined sequence (’N’) were considered (n = 1,169,145) because T STRs without tag may represent false negatives.

**Figure S26 Finding the optimal sequence length for deep learning.** The architecture depicted Supplementary Figure S25 was used to learn features in increasing lengths of sequences centered around T STR ends (x-axis). The accuracy of each model is shown (y-axis).

**Figure S27 Predicting transcription at T STRs with CNN. A.** Accuracy of the deep CNN model was assessed as the Spearman correlation between observed and predicted tag counts of 1,169,145 considered T STRs. Learning was repeated 10 times to ensure model robustness. Results are shown as boxplot (median = 0.81). **B.** Error of one model was assessed as the difference between observed and predicted tag counts. Results are shown for 1,169,145 T repeats as boxplots (median = 1.82).

**Figure S28 Predicting transcription at T repeats.** We built a model in chicken, human’s most distant species considered in FANTOM5 CAGE data. The prediction, though not null, is much less accurate than in human or in mouse (Spearman r = 0.61 and median absolute error = 1.09). Only 58 libraries are available in chicken and the mean signal across these libraries may not be as robust as in human. To confirm that the robustness of the CAGE signal is directly linked to the number of libraries considered, we computed the mean raw tag count in only 58 randomly chosen human libraries and learned a new model using the same architecture as in Supplementary Figure S25. In that case, the signal appears very sparse (bottom) compared to a signal computed on 988 libraries (top) and the model accuracy falls to 0.61. These results reveal that the CAGE signal at T STRs is noisy when considered library-wise but becomes robust when averaged on numerous libraries. Arrows (middle panel) represent T STRs.

**Figure S29 Feature extraction.** A. For each sequence, the effect of changing one nucleotide at a random position into A,T, C or G was assessed. The nucleotide with the maximum predicted tag count is kept and the procedure continues testing another position. This process is repeated over 3,500 randomly selected positions (x-axis) until no further increase of predicted tag count (y-axis) is observed. B. Two sequences were randomly chosen in the test set. The two sequences optimized as in A are then compared position-wise in order to return 1 for each position with the same nucleotide in both sequences and 0 otherwise. This process was repeated for 2,000 pairs of random sequences. y-axis: fraction of identical pairs; x-axis: position (The end of T repeat is located at position 50; repeat of 9 Ts is indicated by red vertical lines). C. A sequence logo was designed from 100 optimized sequences.

**Figure S30 Example of CAGE signal at T STR.** Example of CAGE peak calling from [2] (bottom track) overlapping a 25bp-long T STR (upper track). Significant fraction of the signal is lost when considering an arbitrary window encompassing −20bp/+5bp around T STR end, as indicated by the dashed vertical line. Conversely, this fraction is considered by computing the tag count along STR length.

**Figure S31 Validating the importance of the T STR length. A.** A 9T-long sequence, whose transcription rate is well predicted by the model learned on the arbitrary window (−20bp/+5bp around T repeat end), is chosen (predicted transcription = 6.9967527 / observed transcription = 7). The number of Ts is progressively increased (x-axis) and a prediction is computed for each sequence (y-axis). This reveals that this second model is as sensitive to T repeat length as the model learned on tag count computed along STR length. **B.** The procedure described in Supplementary Figure 29 was repeated with the model learned with a tag count computed on an arbitrary window encompassing 20bp upstream and 5bp downstream T repeat end. A sequence logo was built on 30 optimized sequences revealing that the T repeat length is also learned as a key feature by this second model.

**Figure S32 Motif enrichment comparing T STRs with high and low tag count. A.** Tag count distribution of the top 5,000 sequences with high (blue) and low (red) tag count. **B.** Motif enrichment was computed with HOMER [23] considering 101bp-long sequences centered around T STR ends with high and low tag count (top 5,000). The top12 motifs out of 33 are shown.

**Figure S33 Predicting transcription at T STRs using Random Forest.** The predictive variables are the nucleotide, dinucleotide and trinucleotide content computed into the 50 bp downstream the polyT end and 50 bp upstream the polyT end excluding the T STR. We also integrated the length of the T STR and the nature of the nucleotide following the T STR end (A, C or G at position −1 referring to the CAGE summit at 0). Variable importance was assessed computing the minimal node depth, which reflects the predictiveness of a variable by a depth calculation relative to the root node of a tree (smaller values correspond to more predictive variables). The distribution of minimal depth is represented for top 10 variables according to mean minimal depth. The mean minimal depth was calculated using only non-missing values.

**Figure S34 Distribution of ClinVar variants around permissive CAGE summits.** Position 0 indicates the CAGE summit.

**Figure S35 ClinVar variants induce changes in transcription prediction at T STRs.** Changes were measured as the difference between models scores before and after variant insertion.

## Additional tables

**Table S1 CAGE-associated T STRs are preferentially located in coding genes.** The coordinates of CAGE-associated T STRs (T STR end + 2bp, n = 63,974) and that of all repeats of more than 9 Ts (n = 1,337,561) were intersected with the annotation provided by the FANTOM5 CAGE Associated Transcriptome (CAT). The total number of intersections obtained with CAGE-associated T STRs and all T STRs is 63,480 and 699,048 respectively. The location of all T STRs are shown for comparison. For each gene class, the number of intersections and the corresponding percentage is shown. The median number of CAGE-associated T STRs per gene is 2. In contrast, ~ 47% of all T STRs are located in FANTOM CAT genes with a median number of T STRs per gene of 7. 45,058 out of 63,974 CAGE-associated T STRs are located in introns (*>* 70%), while only 34% of all T STRs are intronic (453,428 out of 1,337,561, Fisher’s exact test p-value *<* 2.2e-16). Likewise, in mouse, 6,500 out of 8,825 CAGE-associated T STRs (~ 74%) but 204,328 out of 834,954 T STRs (~ 24%) are located in introns.

**Table S2 CAGE-associated T-STRs are preferentially located in coding genes.** The coordinates of T STR CAGEs (n = 63,947) and that of all repeats of more than 9 Ts (n = 1,337,561) were intersected with the annotation provided by GENCODEv19. The total number of intersections with T STR CAGEs and all T STRs is 49,312 and 477,492 respectively. The location of all T STRs are shown for comparison. For each gene class, the number of intersections and the corresponding percentage is shown.

**Table S3 FANTOM transcripts and enhancers associated with T STR CAGEs. A.** Start coordinates of polyT-associated FANTOM CAT robust transcripts. The length of the associated polyT tract in the hg19 reference genome is indicated in column 5. **B.** Coordinates of FANTOM enhancer boundaries associated with polyT tracts. The length of the associated polyT tract in the hg19 reference genome is indicated in column 5. Data are provided in a separate excel file.

**Table S4 Predicting transcription at T STRs using LASSO.** The predictive variables are the nucleotide, dinucleotide and trinucleotide content computed into the 50 bp downstream the polyT end and 50 bp upstream the polyT end excluding the T STR. We also integrated the length of the T STR and the nature of the nucleotide following the T STR end (A, C or G at position −1 referring to the CAGE summit at 0). The selected top 10 variables are shown.

**Table S5 Enrichment for disease-associated variant at the vicinity of human T STRs.** The coordinates of ClinVar variants were intersected with that of 101bp-long sequences centered around T STR ends. For each disease, number of intersecting variants is indicated in column 2. For each disease, the fraction of associated variants at T STRs was compared to that of all associated variants using Fisher’s exact test (p-value is indicated in column 3). Data are provided in a separate excel file.

**Table S6 T STR CAGEs in repeat elements.** The coordinates of repetitive elements, according to RepeatMasker, were intersected with that of T STRs associated (n = 63,974) or not (n = 1,273,587) with CAGE peaks. The total number of intersections is indicated in brackets.

**Table S7 Datasets and online resources.**

## References

1. Dunham, I., Kundaje, A., Aldred, S.F., Collins, P.J., Davis, C.A., Doyle, F., Epstein, C.B., Frietze, S., Harrow, J., Kaul, R., Khatun, J., Lajoie, B.R., Landt, S.G., Lee, B.K., Pauli, F., Rosenbloom, K.R., Sabo, P., Safi, A., Sanyal, A., Shoresh, N., Simon, J.M., Song, L., Trinklein, N.D., Altshuler, R.C., Birney, E., Brown, J.B., Cheng, C., Djebali, S., Dong, X., Dunham, I., Ernst, J., Furey, T.S., Gerstein, M., Giardine, B., Greven, M., Hardison, R.C., Harris, R.S., Herrero, J., Hoffman, M.M., Iyer, S., Kellis, M., Khatun, J., Kheradpour, P., Kundaje, A., Lassmann, T., Li, Q., Lin, X., Marinov, G.K., Merkel, A., Mortazavi, A., Parker, S.C., Reddy, T.E., Rozowsky, J., Schlesinger, F., Thurman, R.E., Wang, J., Ward, L.D., Whitfield, T.W., Wilder, S.P., Wu, W., Xi, H.S., Yip, K.Y., Zhuang, J., Pazin, M.J., Lowdon, R.F., Dillon, L.A., Adams, L.B., Kelly, C.J., Zhang, J., Wexler, J.R., Green, E.D., Good, P.J., Feingold, E.A., Bernstein, B.E., Birney, E., Crawford, G.E., Dekker, J., Elnitski, L., Farnham, P.J., Gerstein, M., Giddings, M.C., Gingeras, T.R., Green, E.D., Guigo, R., Hardison, R.C., Hubbard, T.J., Kellis, M., Kent, W., Lieb, J.D., Margulies, E.H., Myers, R.M., Snyder, M., Stamatoyannopoulos, J.A., Tenenbaum, S.A., Weng, Z., White, K.P., Wold, B., Khatun, J., Yu, Y., Wrobel, J., Risk, B.A., Gunawardena, H.P., Kuiper, H.C., Maier, C.W., Xie, L., Chen, X., Giddings, M.C., Bernstein, B.E., Epstein, C.B., Shoresh, N., Ernst, J., Kheradpour, P., Mikkelsen, T.S., Gillespie, S., Goren, A., Ram, O., Zhang, X., Wang, L., Issner, R., Coyne, M.J., Durham, T., Ku, M., Truong, T., Ward, L.D., Altshuler, R.C., Eaton, M.L., Kellis, M., Djebali, S., Davis, C.A., Merkel, A., Dobin, A., Lassmann, T., Mortazavi, A., Tanzer, A., Lagarde, J., Lin, W., Schlesinger, F., Xue, C., Marinov, G.K., Khatun, J., Williams, B.A., Zaleski, C., Rozowsky, J., Roder, M., Kokocinski, F., Abdelhamid, R.F., Alioto, T., Antoshechkin, I., Baer, M.T., Batut, P., Bell, I., Bell, K., Chakrabortty, S., Chen, X., Chrast, J., Curado, J., Derrien, T., Drenkow, J., Dumais, E., Dumais, J., Duttagupta, R., Fastuca, M., Fejes-Toth, K., Ferreira, P., Foissac, S., Fullwood, M.J., Gao, H., Gonzalez, D., Gordon, A., Gunawardena, H.P., Howald, C., Jha, S., Johnson, R., Kapranov, P., King, B., Kingswood, C., Li, G., Luo, O.J., Park, E., Preall, J.B., Presaud, K., Ribeca, P., Risk, B.A., Robyr, D., Ruan, X., Sammeth, M., Sandhu, K.S., Schaeffer, L., See, L.H., Shahab, A., Skancke, J., Suzuki, A.M., Takahashi, H., Tilgner, H., Trout, D., Walters, N., Wang, H., Wrobel, J., Yu, Y., Hayashizaki, Y., Harrow, J., Gerstein, M., Hubbard, T.J., Reymond, A., Antonarakis, S.E., Hannon, G.J., Giddings, M.C., Ruan, Y., Wold, B., Carninci, P., Guigo, R., Gingeras, T.R., Rosenbloom, K.R., Sloan, C.A., Learned, K., Malladi, V.S., Wong, M.C., Barber, G.P., Cline, M.S., Dreszer, T.R., Heitner, S.G., Karolchik, D., Kent, W., Kirkup, V.M., Meyer, L.R., Long, J.C., Maddren, M., Raney, B.J., Furey, T.S., Song, L., Grasfeder, L.L., Giresi, P.G., Lee, B.K., Battenhouse, A., Sheffield, N.C., Simon, J.M., Showers, K.A., Safi, A., London, D., Bhinge, A.A., Shestak, C., Schaner, M.R., Kim, S.K., Zhang, Z.Z., Mieczkowski, P.A., Mieczkowska, J.O., Liu, Z., McDaniell, R.M., Ni, Y., Rashid, N.U., Kim, M.J., Adar, S., Zhang, Z., Wang, T., Winter, D., Keefe, D., Birney, E., Iyer, V.R., Lieb, J.D., Crawford, G.E., Li, G., Sandhu, K.S., Zheng, M., Wang, P., Luo, O.J., Shahab, A., Fullwood, M.J., Ruan, X., Ruan, Y., Myers, R.M., Pauli, F., Williams, B.A., Gertz, J., Marinov, G.K., Reddy, T.E., Vielmetter, J., Partridge, E., Trout, D., Varley, K.E., Gasper, C., Bansal, A., Pepke, S., Jain, P., Amrhein, H., Bowling, K.M., Anaya, M., Cross, M.K., King, B., Muratet, M.A., Antoshechkin, I., Newberry, K.M., McCue, K., Nesmith, A.S., Fisher-Aylor, K.I., Pusey, B., DeSalvo, G., Parker, S.L., Balasubramanian, S., Davis, N.S., Meadows, S.K., Eggleston, T., Gunter, C., Newberry, J., Levy, S.E., Absher, D.M., Mortazavi, A., Wong, W.H., Wold, B., Blow, M.J., Visel, A., Pennachio, L.A., Elnitski, L., Margulies, E.H., Parker, S.C., Petrykowska, H.M., Abyzov, A., Aken, B., Barrell, D., Barson, G., Berry, A., Bignell, A., Boychenko, V., Bussotti, G., Chrast, J., Davidson, C., Derrien, T., Despacio-Reyes, G., Diekhans, M., Ezkurdia, I., Frankish, A., Gilbert, J., Gonzalez, J.M., Griffiths, E., Harte, R., Hendrix, D.A., Howald, C., Hunt, T., Jungreis, I., Kay, M., Khurana, E., Kokocinski, F., Leng, J., Lin, M.F., Loveland, J., Lu, Z., Manthravadi, D., Mariotti, M., Mudge, J., Mukherjee, G., Notredame, C., Pei, B., Rodriguez, J.M., Saunders, G., Sboner, A., Searle, S., Sisu, C., Snow, C., Steward, C., Tanzer, A., Tapanari, E., Tress, M.L., van Baren, M.J., Walters, N., Washietl, S., Wilming, L., Zadissa, A., Zhang, Z., Brent, M., Haussler, D., Kellis, M., Valencia, A., Gerstein, M., Reymond, A., Guigo, R., Harrow, J., Hubbard, T.J., Landt, S.G., Frietze, S., Abyzov, A., Addleman, N., Alexander, R.P., Auerbach, R.K., Balasubramanian, S., Bettinger, K., Bhardwaj, N., Boyle, A.P., Cao, A.R., Cayting, P., Charos, A., Cheng, Y., Cheng, C., Eastman, C., Euskirchen, G., Fleming, J.D., Grubert, F., Habegger, L., Hariharan, M., Harmanci, A., Iyengar, S., Jin, V.X., Karczewski, K.J., Kasowski, M., Lacroute, P., Lam, H., Lamarre-Vincent, N., Leng, J., Lian, J., Lindahl-Allen, M., Min, R., Miotto, B., Monahan, H., Moqtaderi, Z., Mu, X.J., O’Geen, H., Ouyang, Z., Patacsil, D., Pei, B., Raha, D., Ramirez, L., Reed, B., Rozowsky, J., Sboner, A., Shi, M., Sisu, C., Slifer, T., Witt, H., Wu, L., Xu, X., Yan, K.K., Yang, X., Yip, K.Y., Zhang, Z., Struhl, K., Weissman, S.M., Gerstein, M., Farnham, P.J., Snyder, M., Tenenbaum, S.A., Penalva, L.O., Doyle, F., Karmakar, S., Landt, S.G., Bhanvadia, R.R., Choudhury, A., Domanus, M., Ma, L., Moran, J., Patacsil, D., Slifer, T., Victorsen, A., Yang, X., Snyder, M., Auer, T., Centanin, L., Eichenlaub, M., Gruhl, F., Heermann, S., Hoeckendorf, B., Inoue, D., Kellner, T., Kirchmaier, S., Mueller, C., Reinhardt, R., Schertel, L., Schneider, S., Sinn, R., Wittbrodt, B., Wittbrodt, J., Weng, Z., Whitfield, T.W., Wang, J., Collins, P.J., Aldred, S.F., Trinklein, N.D., Partridge, E.C., Myers, R.M., Dekker, J., Jain, G., Lajoie, B.R., Sanyal, A., Balasundaram, G., Bates, D.L., Byron, R., Canfield, T.K., Diegel, M.J., Dunn, D., Ebersol, A.K., Frum, T., Garg, K., Gist, E., Hansen, R., Boatman, L., Haugen, E., Humbert, R., Jain, G., Johnson, A.K., Johnson, E.M., Kutyavin, T.V., Lajoie, B.R., Lee, K., Lotakis, D., Maurano, M.T., Neph, S.J., Neri, F.V., Nguyen, E.D., Qu, H., Reynolds, A.P., Roach, V., Rynes, E., Sabo, P., Sanchez, M.E., Sandstrom, R.S., Sanyal, A., Shafer, A.O., Stergachis, A.B., Thomas, S., Thurman, R.E., Vernot, B., Vierstra, J., Vong, S., Wang, H., Weaver, M.A., Yan, Y., Zhang, M., Akey, J.M., Bender, M., Dorschner, M.O., Groudine, M., MacCoss, M.J., Navas, P., Stamatoyannopoulos, G., Kaul, R., Dekker, J., Stamatoyannopoulos, J.A., Dunham, I., Beal, K., Brazma, A., Flicek, P., Herrero, J., Johnson, N., Keefe, D., Lukk, M., Luscombe, N.M., Sobral, D., Vaquerizas, J.M., Wilder, S.P., Batzoglou, S., Sidow, A., Hussami, N., Kyriazopoulou-Panagiotopoulou, S., Libbrecht, M.W., Schaub, M.A., Kundaje, A., Hardison, R.C., Miller, W., Giardine, B., Harris, R.S., Wu, W., Bickel, P.J., Banfai, B., Boley, N.P., Brown, J.B., Huang, H., Li, Q., Li, J.J., Noble, W.S., Bilmes, J.A., Buske, O.J., Hoffman, M.M., Sahu, A.D., Kharchenko, P.V., Park, P.J., Baker, D., Taylor, J., Weng, Z., Iyer, S., Dong, X., Greven, M., Lin, X., Wang, J., Xi, H.S., Zhuang, J., Gerstein, M., Alexander, R.P., Balasubramanian, S., Cheng, C., Harmanci, A., Lochovsky, L., Min, R., Mu, X.J., Rozowsky, J., Yan, K.K., Yip, K.Y., Birney, E.: An integrated encyclopedia of DNA elements in the human genome. Nature 489(7414), 57–74 (2012)

2. Forrest, A.R., Kawaji, H., Rehli, M., Baillie, J.K., de Hoon, M.J., Haberle, V., Lassmann, T., Kulakovskiy, I.V., Lizio, M., Itoh, M., Andersson, R., Mungall, C.J., Meehan, T.F., Schmeier, S., Bertin, N., J?rgensen, M., Dimont, E., Arner, E., Schmidl, C., Schaefer, U., Medvedeva, Y.A., Plessy, C., Vitezic, M., Severin, J., Semple, C., Ishizu, Y., Young, R.S., Francescatto, M., Alam, I., Albanese, D., Altschuler, G.M., Arakawa, T., Archer, J.A., Arner, P., Babina, M., Rennie, S., Balwierz, P.J., Beckhouse, A.G., Pradhan-Bhatt, S., Blake, J.A., Blumenthal, A., Bodega, B., Bonetti, A., Briggs, J., Brombacher, F., Burroughs, A.M., Califano, A., Cannistraci, C.V., Carbajo, D., Chen, Y., Chierici, M., Ciani, Y., Clevers, H.C., Dalla, E., Davis, C.A., Detmar, M., Diehl, A.D., Dohi, T., Drabl?s, F., Edge, A.S., Edinger, M., Ekwall, K., Endoh, M., Enomoto, H., Fagiolini, M., Fairbairn, L., Fang, H., Farach-Carson, M.C., Faulkner, G.J., Favorov, A.V., Fisher, M.E., Frith, M.C., Fujita, R., Fukuda, S., Furlanello, C., Furino, M., Furusawa, J., Geijtenbeek, T.B., Gibson, A.P., Gingeras, T., Goldowitz, D., Gough, J., Guhl, S., Guler, R., Gustincich, S., Ha, T.J., Hamaguchi, M., Hara, M., Harbers, M., Harshbarger, J., Hasegawa, A., Hasegawa, Y., Hashimoto, T., Herlyn, M., Hitchens, K.J., Ho Sui, S.J., Hofmann, O.M., Hoof, I., Hori, F., Huminiecki, L., Iida, K., Ikawa, T., Jankovic, B.R., Jia, H., Joshi, A., Jurman, G., Kaczkowski, B., Kai, C., Kaida, K., Kaiho, A., Kajiyama, K., Kanamori-Katayama, M., Kasianov, A.S., Kasukawa, T., Katayama, S., Kato, S., Kawaguchi, S., Kawamoto, H., Kawamura, Y.I., Kawashima, T., Kempfle, J.S., Kenna, T.J., Kere, J., Khachigian, L.M., Kitamura, T., Klinken, S.P., Knox, A.J., Kojima, M., Kojima, S., Kondo, N., Koseki, H., Koyasu, S., Krampitz, S., Kubosaki, A., Kwon, A.T., Laros, J.F., Lee, W., Lennartsson, A., Li, K., Lilje, B., Lipovich, L., Mackay-Sim, A., Manabe, R., Mar, J.C., Marchand, B., Mathelier, A., Mejhert, N., Meynert, A., Mizuno, Y., de Lima Morais, D.A., Morikawa, H., Morimoto, M., Moro, K., Motakis, E., Motohashi, H., Mummery, C.L., Murata, M., Nagao-Sato, S., Nakachi, Y., Nakahara, F., Nakamura, T., Nakamura, Y., Nakazato, K., van Nimwegen, E., Ninomiya, N., Nishiyori, H., Noma, S., Noma, S., Noazaki, T., Ogishima, S., Ohkura, N., Ohimiya, H., Ohno, H., Ohshima, M., Okada-Hatakeyama, M., Okazaki, Y., Orlando, V., Ovchinnikov, D.A., Pain, A., Passier, R., Patrikakis, M., Persson, H., Piazza, S., Prendergast, J.G., Rackham, O.J., Ramilowski, J.A., Rashid, M., Ravasi, T., Rizzu, P., Roncador, M., Roy, S., Rye, M.B., Saijyo, E., Sajantila, A., Saka, A., Sakaguchi, S., Sakai, M., Sato, H., Savvi, S., Saxena, A., Schneider, C., Schultes, E.A., Schulze-Tanzil, G.G., Schwegmann, A., Sengstag, T., Sheng, G., Shimoji, H., Shimoni, Y., Shin, J.W., Simon, C., Sugiyama, D., Sugiyama, T., Suzuki, M., Suzuki, N., Swoboda, R.K., ’t Hoen, P.A., Tagami, M., Takahashi, N., Takai, J., Tanaka, H., Tatsukawa, H., Tatum, Z., Thompson, M., Toyodo, H., Toyoda, T., Valen, E., van de Wetering, M., van den Berg, L.M., Verado, R., Vijayan, D., Vorontsov, I.E., Wasserman, W.W., Watanabe, S., Wells, C.A., Winteringham, L.N., Wolvetang, E., Wood, E.J., Yamaguchi, Y., Yamamoto, M., Yoneda, M., Yonekura, Y., Yoshida, S., Zabierowski, S.E., Zhang, P.G., Zhao, X., Zucchelli, S., Summers, K.M., Suzuki, H., Daub, C.O., Kawai, J., Heutink, P., Hide, W., Freeman, T.C., Lenhard, B., Bajic, V.B., Taylor, M.S., Makeev, V.J., Sandelin, A., Hume, D.A., Carninci, P., Hayashizaki, Y.: A promoter-level mammalian expression atlas. Nature 507(7493), 462–470 (2014)

3. Andersson, R., Gebhard, C., Miguel-Escalada, I., Hoof, I., Bornholdt, J., Boyd, M., Chen, Y., Zhao, X., Schmidl, C., Suzuki, T., Ntini, E., Arner, E., Valen, E., Li, K., Schwarzfischer, L., Glatz, D., Raithel, J., Lilje, B., Rapin, N., Bagger, F.O., Jorgensen, M., Andersen, P.R., Bertin, N., Rackham, O., Burroughs, A.M., Baillie, J.K., Ishizu, Y., Shimizu, Y., Furuhata, E., Maeda, S., Negishi, Y., Mungall, C.J., Meehan, T.F., Lassmann, T., Itoh, M., Kawaji, H., Kondo, N., Kawai, J., Lennartsson, A., Daub, C.O., Heutink, P., Hume, D.A., Jensen, T.H., Suzuki, H., Hayashizaki, Y., Muller, F., Forrest, A.R.R., Carninci, P., Rehli, M., Sandelin, A.: An atlas of active enhancers across human cell types and tissues. Nature 507(7493), 455–461 (2014)

4. Hon, C.C., Ramilowski, J.A., Harshbarger, J., Bertin, N., Rackham, O.J., Gough, J., Denisenko, E., Schmeier, S., Poulsen, T.M., Severin, J., Lizio, M., Kawaji, H., Kasukawa, T., Itoh, M., Burroughs, A.M., Noma, S., Djebali, S., Alam, T., Medvedeva, Y.A., Testa, A.C., Lipovich, L., Yip, C.W., Abugessaisa, I., Mendez, M., Hasegawa, A., Tang, D., Lassmann, T., Heutink, P., Babina, M., Wells, C.A., Kojima, S., Nakamura, Y., Suzuki, H., Daub, C.O., de Hoon, M.J., Arner, E., Hayashizaki, Y., Carninci, P., Forrest, A.R.: An atlas of human long non-coding RNAs with accurate 5’ ends. Nature 543(7644), 199–204 (2017)

5. Birney, E., Stamatoyannopoulos, J.A., Dutta, A., Guigo, R., Gingeras, T.R., Margulies, E.H., Weng, Z., Snyder, M., Dermitzakis, E.T., Thurman, R.E., Kuehn, M.S., Taylor, C.M., Neph, S., Koch, C.M., Asthana, S., Malhotra, A., Adzhubei, I., Greenbaum, J.A., Andrews, R.M., Flicek, P., Boyle, P.J., Cao, H., Carter, N.P., Clelland, G.K., Davis, S., Day, N., Dhami, P., Dillon, S.C., Dorschner, M.O., Fiegler, H., Giresi, P.G., Goldy, J., Hawrylycz, M., Haydock, A., Humbert, R., James, K.D., Johnson, B.E., Johnson, E.M., Frum, T.T., Rosenzweig, E.R., Karnani, N., Lee, K., Lefebvre, G.C., Navas, P.A., Neri, F., Parker, S.C., Sabo, P.J., Sandstrom, R., Shafer, A., Vetrie, D., Weaver, M., Wilcox, S., Yu, M., Collins, F.S., Dekker, J., Lieb, J.D., Tullius, T.D., Crawford, G.E., Sunyaev, S., Noble, W.S., Dunham, I., Denoeud, F., Reymond, A., Kapranov, P., Rozowsky, J., Zheng, D., Castelo, R., Frankish, A., Harrow, J., Ghosh, S., Sandelin, A., Hofacker, I.L., Baertsch, R., Keefe, D., Dike, S., Cheng, J., Hirsch, H.A., Sekinger, E.A., Lagarde, J., Abril, J.F., Shahab, A., Flamm, C., Fried, C., Hackermuller, J., Hertel, J., Lindemeyer, M., Missal, K., Tanzer, A., Washietl, S., Korbel, J., Emanuelsson, O., Pedersen, J.S., Holroyd, N., Taylor, R., Swarbreck, D., Matthews, N., Dickson, M.C., Thomas, D.J., Weirauch, M.T., Gilbert, J., Drenkow, J., Bell, I., Zhao, X., Srinivasan, K.G., Sung, W.K., Ooi, H.S., Chiu, K.P., Foissac, S., Alioto, T., Brent, M., Pachter, L., Tress, M.L., Valencia, A., Choo, S.W., Choo, C.Y., Ucla, C., Manzano, C., Wyss, C., Cheung, E., Clark, T.G., Brown, J.B., Ganesh, M., Patel, S., Tammana, H., Chrast, J., Henrichsen, C.N., Kai, C., Kawai, J., Nagalakshmi, U., Wu, J., Lian, Z., Lian, J., Newburger, P., Zhang, X., Bickel, P., Mattick, J.S., Carninci, P., Hayashizaki, Y., Weissman, S., Hubbard, T., Myers, R.M., Rogers, J., Stadler, P.F., Lowe, T.M., Wei, C.L., Ruan, Y., Struhl, K., Gerstein, M., Antonarakis, S.E., Fu, Y., Green, E.D., Karaoz, U., Siepel, A., Taylor, J., Liefer, L.A., Wetterstrand, K.A., Good, P.J., Feingold, E.A., Guyer, M.S., Cooper, G.M., Asimenos, G., Dewey, C.N., Hou, M., Nikolaev, S., Montoya-Burgos, J.I., Loytynoja, A., Whelan, S., Pardi, F., Massingham, T., Huang, H., Zhang, N.R., Holmes, I., Mullikin, J.C., Ureta-Vidal, A., Paten, B., Seringhaus, M., Church, D., Rosenbloom, K., Kent, W.J., Stone, E.A., Batzoglou, S., Goldman, N., Hardison, R.C., Haussler, D., Miller, W., Sidow, A., Trinklein, N.D., Zhang, Z.D., Barrera, L., Stuart, R., King, D.C., Ameur, A., Enroth, S., Bieda, M.C., Kim, J., Bhinge, A.A., Jiang, N., Liu, J., Yao, F., Vega, V.B., Lee, C.W., Ng, P., Shahab, A., Yang, A., Moqtaderi, Z., Zhu, Z., Xu, X., Squazzo, S., Oberley, M.J., Inman, D., Singer, M.A., Richmond, T.A., Munn, K.J., Rada-Iglesias, A., Wallerman, O., Komorowski, J., Fowler, J.C., Couttet, P., Bruce, A.W., Dovey, O.M., Ellis, P.D., Langford, C.F., Nix, D.A., Euskirchen, G., Hartman, S., Urban, A.E., Kraus, P., Van Calcar, S., Heintzman, N., Kim, T.H., Wang, K., Qu, C., Hon, G., Luna, R., Glass, C.K., Rosenfeld, M.G., Aldred, S.F., Cooper, S.J., Halees, A., Lin, J.M., Shulha, H.P., Zhang, X., Xu, M., Haidar, J.N., Yu, Y., Ruan, Y., Iyer, V.R., Green, R.D., Wadelius, C., Farnham, P.J., Ren, B., Harte, R.A., Hinrichs, A.S., Trumbower, H., Clawson, H., Hillman-Jackson, J., Zweig, A.S., Smith, K., Thakkapallayil, A., Barber, G., Kuhn, R.M., Karolchik, D., Armengol, L., Bird, C.P., de Bakker, P.I., Kern, A.D., Lopez-Bigas, N., Martin, J.D., Stranger, B.E., Woodroffe, A., Davydov, E., Dimas, A., Eyras, E., Hallgrimsdottir, I.B., Huppert, J., Zody, M.C., Abecasis, G.R., Estivill, X., Bouffard, G.G., Guan, X., Hansen, N.F., Idol, J.R., Maduro, V.V., Maskeri, B., McDowell, J.C., Park, M., Thomas, P.J., Young, A.C., Blakesley, R.W., Muzny, D.M., Sodergren, E., Wheeler, D.A., Worley, K.C., Jiang, H., Weinstock, G.M., Gibbs, R.A., Graves, T., Fulton, R., Mardis, E.R., Wilson, R.K., Clamp, M., Cuff, J., Gnerre, S., Jaffe, D.B., Chang, J.L., Lindblad-Toh, K., Lander, E.S., Koriabine, M., Nefedov, M., Osoegawa, K., Yoshinaga, Y., Zhu, B., de Jong, P.J.: Identification and analysis of functional elements in 1human genome by the ENCODE pilot project. Nature 447(7146), 799–816 (2007)

6. Carninci, P., Kasukawa, T., Katayama, S., Gough, J., Frith, M.C., Maeda, N., Oyama, R., Ravasi, T., Lenhard, B., Wells, C., Kodzius, R., Shimokawa, K., Bajic, V.B., Brenner, S.E., Batalov, S., Forrest, A.R., Zavolan, M., Davis, M.J., Wilming, L.G., Aidinis, V., Allen, J.E., Ambesi-Impiombato, A., Apweiler, R., Aturaliya, R.N., Bailey, T.L., Bansal, M., Baxter, L., Beisel, K.W., Bersano, T., Bono, H., Chalk, A.M., Chiu, K.P., Choudhary, V., Christoffels, A., Clutterbuck, D.R., Crowe, M.L., Dalla, E., Dalrymple, B.P., de Bono, B., Della Gatta, G., di Bernardo, D., Down, T., Engstrom, P., Fagiolini, M., Faulkner, G., Fletcher, C.F., Fukushima, T., Furuno, M., Futaki, S., Gariboldi, M., Georgii-Hemming, P., Gingeras, T.R., Gojobori, T., Green, R.E., Gustincich, S., Harbers, M., Hayashi, Y., Hensch, T.K., Hirokawa, N., Hill, D., Huminiecki, L., Iacono, M., Ikeo, K., Iwama, A., Ishikawa, T., Jakt, M., Kanapin, A., Katoh, M., Kawasawa, Y., Kelso, J., Kitamura, H., Kitano, H., Kollias, G., Krishnan, S.P., Kruger, A., Kummerfeld, S.K., Kurochkin, I.V., Lareau, L.F., Lazarevic, D., Lipovich, L., Liu, J., Liuni, S., McWilliam, S., Madan Babu, M., Madera, M., Marchionni, L., Matsuda, H., Matsuzawa, S., Miki, H., Mignone, F., Miyake, S., Morris, K., Mottagui-Tabar, S., Mulder, N., Nakano, N., Nakauchi, H., Ng, P., Nilsson, R., Nishiguchi, S., Nishikawa, S., Nori, F., Ohara, O., Okazaki, Y., Orlando, V., Pang, K.C., Pavan, W.J., Pavesi, G., Pesole, G., Petrovsky, N., Piazza, S., Reed, J., Reid, J.F., Ring, B.Z., Ringwald, M., Rost, B., Ruan, Y., Salzberg, S.L., Sandelin, A., Schneider, C., Schonbach, C., Sekiguchi, K., Semple, C.A., Seno, S., Sessa, L., Sheng, Y., Shibata, Y., Shimada, H., Shimada, K., Silva, D., Sinclair, B., Sperling, S., Stupka, E., Sugiura, K., Sultana, R., Takenaka, Y., Taki, K., Tammoja, K., Tan, S.L., Tang, S., Taylor, M.S., Tegner, J., Teichmann, S.A., Ueda, H.R., van Nimwegen, E., Verardo, R., Wei, C.L., Yagi, K., Yamanishi, H., Zabarovsky, E., Zhu, S., Zimmer, A., Hide, W., Bult, C., Grimmond, S.M., Teasdale, R.D., Liu, E.T., Brusic, V., Quackenbush, J., Wahlestedt, C., Mattick, J.S., Hume, D.A., Kai, C., Sasaki, D., Tomaru, Y., Fukuda, S., Kanamori-Katayama, M., Suzuki, M., Aoki, J., Arakawa, T., Iida, J., Imamura, K., Itoh, M., Kato, T., Kawaji, H., Kawagashira, N., Kawashima, T., Kojima, M., Kondo, S., Konno, H., Nakano, K., Ninomiya, N., Nishio, T., Okada, M., Plessy, C., Shibata, K., Shiraki, T., Suzuki, S., Tagami, M., Waki, K., Watahiki, A., Okamura-Oho, Y., Suzuki, H., Kawai, J., Hayashizaki, Y.: The transcriptional landscape of the mammalian genome. Science 309(5740), 1559–1563 (2005)

7. Ard, R., Allshire, R.C., Marquardt, S.: Emerging Properties and Functional Consequences of Noncoding Transcription. Genetics 207(2), 357–367 (2017)

8. Palazzo, A.F., Lee, E.S.: Non-coding RNA: what is functional and what is junk? Front Genet 6, 2 (2015)

9. Cheneby, J., Gheorghe, M., Artufel, M., Mathelier, A., Ballester, B.: ReMap 2018: an updated atlas of regulatory regions from an integrative analysis of DNA-binding ChIP-seq experiments. Nucleic Acids Res. (2017)

10. Schaub, M.A., Boyle, A.P., Kundaje, A., Batzoglou, S., Snyder, M.: Linking disease associations with regulatory information in the human genome. Genome Res. 22(9), 1748–1759 (2012)

11. Maurano, M.T., Humbert, R., Rynes, E., Thurman, R.E., Haugen, E., Wang, H., Reynolds, A.P., Sandstrom, R., Qu, H., Brody, J., Shafer, A., Neri, F., Lee, K., Kutyavin, T., Stehling-Sun, S., Johnson, A.K., Canfield, T.K., Giste, E., Diegel, M., Bates, D., Hansen, R.S., Neph, S., Sabo, P.J., Heimfeld, S., Raubitschek, A., Ziegler, S., Cotsapas, C., Sotoodehnia, N., Glass, I., Sunyaev, S.R., Kaul, R., Stamatoyannopoulos, J.A.: Systematic localization of common disease-associated variation in regulatory DNA. Science 337(6099), 1190–1195 (2012)

12. Kellis, M., Wold, B., Snyder, M.P., Bernstein, B.E., Kundaje, A., Marinov, G.K., Ward, L.D., Birney, E., Crawford, G.E., Dekker, J., Dunham, I., Elnitski, L.L., Farnham, P.J., Feingold, E.A., Gerstein, M., Giddings, M.C., Gilbert, D.M., Gingeras, T.R., Green, E.D., Guigo, R., Hubbard, T., Kent, J., Lieb, J.D., Myers, R.M., Pazin, M.J., Ren, B., Stamatoyannopoulos, J.A., Weng, Z., White, K.P., Hardison, R.C.: Defining functional DNA elements in the human genome. Proc. Natl. Acad. Sci. U.S.A. 111(17), 6131–6138 (2014)

13. Clark, M.B., Choudhary, A., Smith, M.A., Taft, R.J., Mattick, J.S.: The dark matter rises: the expanding world of regulatory RNAs. Essays Biochem. 54, 1–16 (2013)

14. Willems, T., Gymrek, M., Highnam, G., Mittelman, D., Erlich, Y.: The landscape of human STR variation. Genome Res. 24(11), 1894–1904 (2014)

15. Gymrek, M., Willems, T., Guilmatre, A., Zeng, H., Markus, B., Georgiev, S., Daly, M.J., Price, A.L., Pritchard, J.K., Sharp, A.J., Erlich, Y.: Abundant contribution of short tandem repeats to gene expression variation in humans. Nat. Genet. 48(1), 22–29 (2016)

16. Press, M.O., McCoy, R.C., Hall, A.N., Akey, J.M., Queitsch, C.: Massive variation of short tandem repeats with functional consequences across strains of Arabidopsis thaliana. Genome Res. 28(8), 1169–1178 (2018)

17. Rothenburg, S., Koch-Nolte, F., Rich, A., Haag, F.: A polymorphic dinucleotide repeat in the rat nucleolin gene forms Z-DNA and inhibits promoter activity. Proc. Natl. Acad. Sci. U.S.A. 98(16), 8985–8990 (2001)

18. Contente, A., Dittmer, A., Koch, M.C., Roth, J., Dobbelstein, M.: A polymorphic microsatellite that mediates induction of PIG3 by p53. Nat. Genet. 30(3), 315–320 (2002)

19. Martin, P., Makepeace, K., Hill, S.A., Hood, D.W., Moxon, E.R.: Microsatellite instability regulates transcription factor binding and gene expression. Proc. Natl. Acad. Sci. U.S.A. 102(10), 3800–3804 (2005)

20. Kanamori-Katayama, M., Itoh, M., Kawaji, H., Lassmann, T., Katayama, S., Kojima, M., Bertin, N., Kaiho, A., Ninomiya, N., Daub, C.O., Carninci, P., Forrest, A.R., Hayashizaki, Y.: Unamplified cap analysis of gene expression on a single-molecule sequencer. Genome Res. 21(7), 1150–1159 (2011)

21. Murata, M., Nishiyori-Sueki, H., Kojima-Ishiyama, M., Carninci, P., Hayashizaki, Y., Itoh, M.: Detecting expressed genes using CAGE. Methods Mol. Biol. 1164, 67–85 (2014)

22. Li, C., Lenhard, B., Luscombe, N.M.: Integrated analysis sheds light on evolutionary trajectories of young transcription start sites in the human genome. Genome Res. 28(5), 676–688 (2018)

23. Heinz, S., Benner, C., Spann, N., Bertolino, E., Lin, Y.C., Laslo, P., Cheng, J.X., Murre, C., Singh, H., Glass, C.K.: Simple combinations of lineage-determining transcription factors prime cis-regulatory elements required for macrophage and B cell identities. Mol. Cell 38(4), 576–589 (2010)

24. Smale, S.T., Baltimore, D.: The ”initiator” as a transcription control element. Cell 57(1), 103–113 (1989)

25. Butler, J.E., Kadonaga, J.T.: The RNA polymerase II core promoter: a key component in the regulation of gene expression. Genes Dev. 16(20), 2583–2592 (2002)

26. Dechering, K.J., Cuelenaere, K., Konings, R.N., Leunissen, J.A.: Distinct frequency-distributions of homopolymeric DNA tracts in different genomes. Nucleic Acids Res. 26(17), 4056–4062 (1998)

27. Scruggs, B.S., Gilchrist, D.A., Nechaev, S., Muse, G.W., Burkholder, A., Fargo, D.C., Adelman, K.: Bidirectional Transcription Arises from Two Distinct Hubs of Transcription Factor Binding and Active Chromatin. Mol. Cell 58(6), 1101–1112 (2015)

28. Neri, F., Rapelli, S., Krepelova, A., Incarnato, D., Parlato, C., Basile, G., Maldotti, M., Anselmi, F., Oliviero, S.: Intragenic DNA methylation prevents spurious transcription initiation. Nature 543(7643), 72–77 (2017)

29. Nielsen, M., Ard, R., Leng, X., Ivanov, M., Kindgren, P., Pelechano, V., Marquardt, S.: Transcription-driven Chromatin repression of Intragenic transcription start sites. PLoS Genet. 15(2), 1007969 (2019)

30. Andersson, R., Refsing Andersen, P., Valen, E., Core, L.J., Bornholdt, J., Boyd, M., Heick Jensen, T., Sandelin, A.: Nuclear stability and transcriptional directionality separate functionally distinct RNA species. Nat Commun 5, 5336 (2014)

31. de Rie, D., Abugessaisa, I., Alam, T., Arner, E., Arner, P., Ashoor, H., Astrom, G., Babina, M., Bertin, N., Burroughs, A.M., Carlisle, A.J., Daub, C.O., Detmar, M., Deviatiiarov, R., Fort, A., Gebhard, C., Goldowitz, D., Guhl, S., Ha, T.J., Harshbarger, J., Hasegawa, A., Hashimoto, K., Herlyn, M., Heutink, P., Hitchens, K.J., Hon, C.C., Huang, E., Ishizu, Y., Kai, C., Kasukawa, T., Klinken, P., Lassmann, T., Lecellier, C.H., Lee, W., Lizio, M., Makeev, V., Mathelier, A., Medvedeva, Y.A., Mejhert, N., Mungall, C.J., Noma, S., Ohshima, M., Okada-Hatakeyama, M., Persson, H., Rizzu, P., Roudnicky, F., S?trom, P., Sato, H., Severin, J., Shin, J.W., Swoboda, R.K., Tarui, H., Toyoda, H., Vitting-Seerup, K., Winteringham, L., Yamaguchi, Y., Yasuzawa, K., Yoneda, M., Yumoto, N., Zabierowski, S., Zhang, P.G., Wells, C.A., Summers, K.M., Kawaji, H., Sandelin, A., Rehli, M., Hayashizaki, Y., Carninci, P., Forrest, A.R.R., de Hoon, M.J.L.: An integrated expression atlas of miRNAs and their promoters in human and mouse. Nat. Biotechnol. 35(9), 872–878 (2017)

32. Corces, M.R., Granja, J.M., Shams, S., Louie, B.H., Seoane, J.A., Zhou, W., Silva, T.C., Groeneveld, C., Wong, C.K., Cho, S.W., Satpathy, A.T., Mumbach, M.R., Hoadley, K.A., Robertson, A.G., Sheffield, N.C., Felau, I., Castro, M.A.A., Berman, B.P., Staudt, L.M., Zenklusen, J.C., Laird, P.W., Curtis, C., Greenleaf, W.J., Chang, H.Y., Akbani, R., Benz, C.C., Boyle, E.A., Broom, B.M., Cherniack, A.D., Craft, B., Demchok, J.A., Doane, A.S., Elemento, O., Ferguson, M.L., Goldman, M.J., Hayes, D.N., He, J., Hinoue, T., Imielinski, M., Jones, S.J.M., Kemal, A., Knijnenburg, T.A., Korkut, A., Lin, D.C., Liu, Y., Mensah, M.K.A., Mills, G.B., Reuter, V.P., Schultz, A., Shen, H., Smith, J.P., Tarnuzzer, R., Trefflich, S., Wang, Z., Weinstein, J.N., Westlake, L.C., Xu, J., Yang, L., Yau, C., Zhao, Y., Zhu, J.: The chromatin accessibility landscape of primary human cancers. Science 362(6413) (2018)

33. Ernst, J., Kellis, M.: ChromHMM: automating chromatin-state discovery and characterization. Nature Methods 9(3), 215–216 (2012). doi:10.1038/nmeth.1906. Accessed 2016-07-15

34. Hoffman, M.M., Buske, O.J., Wang, J., Weng, Z., Bilmes, J.A., Noble, W.S.: Unsupervised pattern discovery in human chromatin structure through genomic segmentation. Nature Methods 9(5), 473–476 (2012). doi:10.1038/nmeth.1937. Accessed 2016-07-15

35. Natoli, G., Andrau, J.C.: Noncoding transcription at enhancers: general principles and functional models. Annu. Rev. Genet. 46, 1–19 (2012)

36. Guttman, M., Amit, I., Garber, M., French, C., Lin, M.F., Feldser, D., Huarte, M., Zuk, O., Carey, B.W., Cassady, J.P., Cabili, M.N., Jaenisch, R., Mikkelsen, T.S., Jacks, T., Hacohen, N., Bernstein, B.E., Kellis, M., Regev, A., Rinn, J.L., Lander, E.S.: Chromatin signature reveals over a thousand highly conserved large non-coding RNAs in mammals. Nature 458(7235), 223–227 (2009)

37. Pekowska, A., Benoukraf, T., Ferrier, P., Spicuglia, S.: A unique H3K4me2 profile marks tissue-specific gene regulation. Genome Res. 20(11), 1493–1502 (2010)

38. Wang, Y., Li, X., Hu, H.: H3K4me2 reliably defines transcription factor binding regions in different cells. Genomics 103(2-3), 222–228 (2014)

39. Li, G., Ruan, X., Auerbach, R.K., Sandhu, K.S., Zheng, M., Wang, P., Poh, H.M., Goh, Y., Lim, J., Zhang, J., Sim, H.S., Peh, S.Q., Mulawadi, F.H., Ong, C.T., Orlov, Y.L., Hong, S., Zhang, Z., Landt, S., Raha, D., Euskirchen, G., Wei, C.L., Ge, W., Wang, H., Davis, C., Fisher-Aylor, K.I., Mortazavi, A., Gerstein, M., Gingeras, T., Wold, B., Sun, Y., Fullwood, M.J., Cheung, E., Liu, E., Sung, W.K., Snyder, M., Ruan, Y.: Extensive promoter-centered chromatin interactions provide a topological basis for transcription regulation. Cell 148(1-2), 84–98 (2012)

40. Eraslan, G., Avsec, Z., Gagneur, J., Theis, F.J.: Deep learning: new computational modelling techniques for genomics. Nat. Rev. Genet. 20(7), 389–403 (2019)

41. Tibshirani, R.: Regression shrinkage and selection via the lasso. Journal of the Royal Statistical Society. Series B (Methodological), 267–288 (1996)

42. Liaw, A., Wiener, M.: Classification and regression by randomforest. R News 2(3), 18–22 (2002)

43. Payton, A., Sindrewicz, P., Pessoa, V., Platt, H., Horan, M., Ollier, W., Bubb, V.J., Pendleton, N., Quinn, J.P.: A TOMM40 poly-T variant modulates gene expression and is associated with vocabulary ability and decline in nonpathologic aging. Neurobiol. Aging 39, 1–7 (2016)

44. Linnertz, C., Anderson, L., Gottschalk, W., Crenshaw, D., Lutz, M.W., Allen, J., Saith, S., Mihovilovic, M., Burke, J.R., Welsh-Bohmer, K.A., Roses, A.D., Chiba-Falek, O.: The cis-regulatory effect of an Alzheimer’s disease-associated poly-T locus on expression of TOMM40 and apolipoprotein E genes. Alzheimers Dement 10(5), 541–551 (2014)

45. Maruszak, A., Peplonska, B., Safranow, K., Chodakowska-Zebrowska, M., Barcikowska, M., Zekanowski, C.: TOMM40 rs10524523 polymorphism’s role in late-onset Alzheimer’s disease and in longevity. J. Alzheimers Dis. 28(2), 309–322 (2012)

46. Greenbaum, L., Springer, R.R., Lutz, M.W., Heymann, A., Lubitz, I., Cooper, I., Kravitz, E., Sano, M., Roses, A.D., Silverman, J.M., Saunders, A.M., Beeri, M.S.: The TOMM40 poly-T rs10524523 variant is associated with cognitive performance among non-demented elderly with type 2 diabetes. Eur Neuropsychopharmacol 24(9), 1492–1499 (2014)

47. Bernardi, L., Gallo, M., Anfossi, M., Conidi, M.E., Colao, R., Puccio, G., Curcio, S.A., Frangipane, F., Clodomiro, A., Mirabelli, M., Vasso, F., Smirne, N., Di Lorenzo, R., Maletta, R., Bruni, A.C.: Role of TOMM40 rs10524523 polymorphism in onset of alzheimer’s disease caused by the PSEN1 M146L mutation. J. Alzheimers Dis. 37(2), 285–289 (2013)

48. Mastaglia, F.L., Rojana-udomsart, A., James, I., Needham, M., Day, T.J., Kiers, L., Corbett, J.A., Saunders, A.M., Lutz, M.W., Roses, A.D.: Polymorphism in the TOMM40 gene modifies the risk of developing sporadic inclusion body myositis and the age of onset of symptoms. Neuromuscul. Disord. 23(12), 969–974 (2013)

49. Zhao, L.J., Zhang, S., Chinnadurai, G.: Sox9 transactivation and testicular expression of a novel human gene, KIAA0800. J. Cell. Biochem. 86(2), 277–289 (2002)

50. Feupe Fotsing, S., Wang, C., Saini, S., Shleizer-Burko, S., Goren, A., Gymrek, M.: Multi-tissue analysis reveals short tandem repeats as ubiquitous regulators of gene expression and complex traits. bioRxiv (2018). doi:10.1101/495226. https://www.biorxiv.org/content/early/2018/12/13/495226.full.pdf

51. Sherry, S.T., Ward, M.H., Kholodov, M., Baker, J., Phan, L., Smigielski, E.M., Sirotkin, K.: dbSNP: the NCBI database of genetic variation. Nucleic Acids Res. 29(1), 308–311 (2001)

52. Landrum, M.J., Lee, J.M., Benson, M., Brown, G., Chao, C., Chitipiralla, S., Gu, B., Hart, J., Hoffman, D., Hoover, J., Jang, W., Katz, K., Ovetsky, M., Riley, G., Sethi, A., Tully, R., Villamarin-Salomon, R., Rubinstein, W., Maglott, D.R.: ClinVar: public archive of interpretations of clinically relevant variants. Nucleic Acids Res. 44(D1), 862–868 (2016)

53. Core, L.J., Martins, A.L., Danko, C.G., Waters, C.T., Siepel, A., Lis, J.T.: Analysis of nascent RNA identifies a unified architecture of initiation regions at mammalian promoters and enhancers. Nat. Genet. 46(12), 1311–1320 (2014)

54. Bergmann, J.H., Li, J., Eckersley-Maslin, M.A., Rigo, F., Freier, S.M., Spector, D.L.: Regulation of the ESC transcriptome by nuclear long noncoding RNAs. Genome Res. 25(9), 1336–1346 (2015)

55. Derrien, T., Johnson, R., Bussotti, G., Tanzer, A., Djebali, S., Tilgner, H., Guernec, G., Martin, D., Merkel, A., Knowles, D.G., Lagarde, J., Veeravalli, L., Ruan, X., Ruan, Y., Lassmann, T., Carninci, P., Brown, J.B., Lipovich, L., Gonzalez, J.M., Thomas, M., Davis, C.A., Shiekhattar, R., Gingeras, T.R., Hubbard, T.J., Notredame, C., Harrow, J., Guigo, R.: The GENCODE v7 catalog of human long noncoding RNAs: analysis of their gene structure, evolution, and expression. Genome Res. 22(9), 1775–1789 (2012)

56. Djebali, S., Davis, C.A., Merkel, A., Dobin, A., Lassmann, T., Mortazavi, A., Tanzer, A., Lagarde, J., Lin, W., Schlesinger, F., Xue, C., Marinov, G.K., Khatun, J., Williams, B.A., Zaleski, C., Rozowsky, J., Roder, M., Kokocinski, F., Abdelhamid, R.F., Alioto, T., Antoshechkin, I., Baer, M.T., Bar, N.S., Batut, P., Bell, K., Bell, I., Chakrabortty, S., Chen, X., Chrast, J., Curado, J., Derrien, T., Drenkow, J., Dumais, E., Dumais, J., Duttagupta, R., Falconnet, E., Fastuca, M., Fejes-Toth, K., Ferreira, P., Foissac, S., Fullwood, M.J., Gao, H., Gonzalez, D., Gordon, A., Gunawardena, H., Howald, C., Jha, S., Johnson, R., Kapranov, P., King, B., Kingswood, C., Luo, O.J., Park, E., Persaud, K., Preall, J.B., Ribeca, P., Risk, B., Robyr, D., Sammeth, M., Schaffer, L., See, L.H., Shahab, A., Skancke, J., Suzuki, A.M., Takahashi, H., Tilgner, H., Trout, D., Walters, N., Wang, H., Wrobel, J., Yu, Y., Ruan, X., Hayashizaki, Y., Harrow, J., Gerstein, M., Hubbard, T., Reymond, A., Antonarakis, S.E., Hannon, G., Giddings, M.C., Ruan, Y., Wold, B., Carninci, P., Guigo, R., Gingeras, T.R.: Landscape of transcription in human cells. Nature 489(7414), 101–108 (2012)

57. Matylla-Kulinska, K., Tafer, H., Weiss, A., Schroeder, R.: Functional repeat-derived RNAs often originate from retrotransposon-propagated ncRNAs. Wiley Interdiscip Rev RNA 5(5), 591–600 (2014)

58. Ferreira, D., Meles, S., Escudeiro, A., Mendes-da-Silva, A., Adega, F., Chaves, R.: Satellite non-coding RNAs: the emerging players in cells, cellular pathways and cancer. Chromosome Res. 23(3), 479–493 (2015)

59. Goodier, J.L.: Restricting retrotransposons: a review. Mob DNA 7, 16 (2016)

60. Gao, B., Wang, S., Wang, Y., Shen, D., Xue, S., Chen, C., Cui, H., Song, C.: Low diversity, activity, and density of transposable elements in five avian genomes. Funct. Integr. Genomics 17(4), 427–439 (2017)

61. Roy-Engel, A.M., Salem, A.H., Oyeniran, O.O., Deininger, L., Hedges, D.J., Kilroy, G.E., Batzer, M.A., Deininger, P.L.: Active Alu element ”A-tails”: size does matter. Genome Res. 12(9), 1333–1344 (2002)

62. Dewannieux, M., Heidmann, T.: Role of poly(A) tail length in Alu retrotransposition. Genomics 86(3), 378–381 (2005)

63. Segal, E., Widom, J.: Poly(dA:dT) tracts: major determinants of nucleosome organization. Curr. Opin. Struct. Biol. 19(1), 65–71 (2009)

64. Krietenstein, N., Wal, M., Watanabe, S., Park, B., Peterson, C.L., Pugh, B.F., Korber, P.: Genomic Nucleosome Organization Reconstituted with Pure Proteins. Cell 167(3), 709–721 (2016)

65. Weingarten-Gabbay, S., Nir, R., Lubliner, S., Sharon, E., Kalma, Y., Weinberger, A., Segal, E.: Systematic interrogation of human promoters. Genome Res. 29(2), 171–183 (2019)

66. Martin, D., Maillol, V., Rivals, E.: Fast and accurate genome-scale identification of DNA-binding sites. In: BIBM: Bioinformatics and Biomedicine, Madrid, Spain, pp. 201–205 (2018). This is the author version of the article published in the conference proceedings. It includes supplementary information. A software called MOTIF is available on the ATGC bioinformatics platform. https://hal-lirmm.ccsd.cnrs.fr/lirmm-01967466

67. Quinlan, A.R., Hall, I.M.: BEDTools: a flexible suite of utilities for comparing genomic features. Bioinformatics (Oxford, England) 26(6), 841–842 (2010). doi:10.1093/bioinformatics/btq033

68. Pohl, A., Beato, M.: bwtool: a tool for bigWig files. Bioinformatics 30(11), 1618–1619 (2014). doi:10.1093/bioinformatics/btu056. Accessed 2015-07-10

69. Ramirez, F., Ryan, D.P., Gruning, B., Bhardwaj, V., Kilpert, F., Richter, A.S., Heyne, S., Dundar, F., Manke, T.: deepTools2: a next generation web server for deep-sequencing data analysis. Nucleic Acids Res. 44(W1), 160–165 (2016)

70. R Core Team: R: A Language and Environment for Statistical Computing. R Foundation for Statistical Computing, Vienna, Austria (2017). R Foundation for Statistical Computing. https://www.R-project.org/

71. Wickham, H.: Ggplot2: Elegant Graphics for Data Analysis. Springer, ??? (2016). http://ggplot2.org

72. Li, H., Handsaker, B., Wysoker, A., Fennell, T., Ruan, J., Homer, N., Marth, G., Abecasis, G., Durbin, R.: The Sequence Alignment/Map format and SAMtools. Bioinformatics 25(16), 2078–2079 (2009)

73. Severin, J., Lizio, M., Harshbarger, J., Kawaji, H., Daub, C.O., Hayashizaki, Y., Bertin, N., Forrest, A.R.: Interactive visualization and analysis of large-scale sequencing datasets using ZENBU. Nat. Biotechnol. 32(3), 217–219 (2014)

74. Dalke, A., Wilczynski, B., Chapman, B.A., Cox, C.J., Kauff, F., Friedberg, I., Chang, J.T., de Hoon, M.J.L., Cock, P.J.A., Hamelryck, T., Antao, T.: Biopython: freely available Python tools for computational molecular biology and bioinformatics. Bioinformatics 25(11), 1422–1423 (2009). doi:10.1093/bioinformatics/btp163. http://oup.prod.sis.lan/bioinformatics/article-pdf/25/11/1422/944180/btp163.pdf

75. Chollet, F., et al.: Keras. https://keras.io (2015)

76. Abadi, M., Agarwal, A., Barham, P., Brevdo, E., Chen, Z., Citro, C., Corrado, G.S., Davis, A., Dean, J., Devin, M., Ghemawat, S., Goodfellow, I., Harp, A., Irving, G., Isard, M., Jia, Y., Jozefowicz, R., Kaiser, L., Kudlur, M., Levenberg, J., Mané, D., Monga, R., Moore, S., Murray, D., Olah, C., Schuster, M., Shlens, J., Steiner, B., Sutskever, I., Talwar, K., Tucker, P., Vanhoucke, V., Vasudevan, V., Viégas, F., Vinyals, O., Warden, P., Wattenberg, M., Wicke, M., Yu, Y., Zheng, X.: TensorFlow: Large-Scale Machine Learning on Heterogeneous Systems. Software available from tensorflow.org (2015). http://tensorflow.org/

77. McKinney, W.: Data structures for statistical computing in python. In: van der Walt, S., Millman, J. (eds.) Proceedings of the 9th Python in Science Conference, pp. 51–56 (2010)

78. Wei, W., Pelechano, V., Jarvelin, A.I., Steinmetz, L.M.: Functional consequences of bidirectional promoters. Trends Genet. 27(7), 267–276 (2011)

